# Purinoreceptors and ectonucleotidases control ATP-induced calcium waveforms and calcium-dependent responses in microglia

**DOI:** 10.1101/2021.06.19.448892

**Authors:** Byeong Jae Chun, Surya Aryal, Bin Sun, Josh Bruno, Chris Richards, Adam D. Bachstetter, Peter M. Kekenes-Huskey

**Affiliations:** Department of Cell and Molecular Physiology, Loyola University Chicago; Department of Chemistry, University of Kentucky; College of Medicine, University of Kentucky

## Abstract

Adenosine triphosphate (ATP) drives microglia motility and cytokine production by activating P2X- and P2Y- class purinergic receptors with extracellular ATP and its metabolites. Purinergic receptor activation gives rise to diverse intracellular Ca^2+^ signals, or waveforms, that differ in amplitude, duration, and frequency. Whether and how these diverse waveforms influence microglia function is not well established. We developed a computational model trained with published primary murine microglia studies. We simulate how purinoreceptors influence Ca^2+^ signaling and migration and how purinoreceptor expression modifies these processes. Our simulation confirmed that P2 receptors encode the amplitude and duration of the ATP-induced calcium waveforms. Our simulations also implicate CD39, an ectonucleotidase that rapidly degrades ATP, as a regulator of purinergic receptor-induced Ca^2+^ responses. We, therefore, next evaluated how purinoreceptors and ectonucleotidase work in tandem. Our modeling results indicate that small transients are sufficient to promote motility, while large and sustained transients are needed for cytokine responses. Lastly, we predict how these phenotypical responses vary in a BV2 microglia cell line using published P2 receptor mRNA data to illustrate how our computer model can be extrapolated to diverse microglia subtypes. These findings provide important insights into how differences in purinergic receptor expression influence the microglia’s responses to ATP.

## 2 Introduction

Microglia are the central nervous system (CNS) macrophages. They contribute homeostatic and innate immune responses when subject to a spectrum of molecular stimuli, including those associated with infection and cellular damage. Microglia respond to these stimuli by migrating, undergoing changes in gene programming, secreting cytokines and chemokines to engage the adaptive immune response, and phagocytosing foreign bodies. Many details of these complex signaling pathways controlling microglial responses to such cues are beginning to emerge, including those mediated by adenosine triphosphate (ATP) and its nucleotide derivatives (45).

Extracellular ATP invokes Ca^2+^ waveforms in microglia that trigger or influence cytokine and motility responses (59; 35; 33), as well as a broad set of microglial signaling pathways (42; 56). In other cell types, the waveform of an induced Ca^2+^ signal, that is, its duration, amplitude and frequency, has been shown to selectively control intracellular processes including phosphorylation, gene transcription and mechanical responses (11). Despite these observations, it has yet to be determined if the properties of Ca^2+^ waveforms in microglia exhibit similar selective control of their physiological functions.

ATP-dependent responses in microglia are mediated by purinergic (P2) receptors. P2 receptors are broadly categorized into two classes: ionotropic (P2X) and metabotropic (P2Y) receptors. Ionotropic P2X receptor (*P*2*X*) receptors are non-selective cation channels widely expressed in cells throughout the CNS including microglia. Of these, *P*2*X*7 and *P*2*X*4 tend to be the most highly expressed *P*2*X*-class receptors in microglia (42; 2). In our previous work (9), we developed a computational model demonstrating that *P*2*X* activation promotes the production of a pro-inflammatory cytokine, tumor necrosis factor alpha (TNF*α*). However, microglia also express P2Y receptor (*P*2*Y*) receptors that comprise G protein coupled receptors (GPCR), which mediate pathways including endoplasmic reticulum (ER) Ca^2+^ release (42; 56). There are several prominent *P*2*Y* receptors present in microglia such as P2Y2, P2Y6, P2Y12, and P2Y13 (32). In principle, both classes of P2 receptors contribute to ATP-mediated responses in microglia, but their cocurrent contributions have yet to be determined in quantitative detail.

Microglia exhibit changes in motility, e.g. the extension and retraction of plasma membrane, in response to ATP and can migrate toward sources of ATP. Ca^2+^ transients that coincide with directed motility and migration have been observed in microglia (58; 59; 35; 24). Given the diverse Ca^2+^ waveforms induced by P2 activation, there is an intriguing possibility that microglia adopt unique cell responses to different waveforms that could select for migration versus inflammatory behaviors. However, it remains to be determined if variable Ca^2+^ waveforms are just a consequence of ATP stimulation or if they selectively influence cell funcions.

Ca^2+^ waveforms and the capacity for cell migration in response to ATP are dependent on P2 expression and activity. P2 subtype expression can vary considerably among microglial subpopulations and activation states (30; 15; 5). As an example, resting, *in vivo* microglia are characterized by having high P2Y12 receptor (*P*2*Y* 12) expression and comparatively low expression of *P*2*X*4 and *P*2*X*7 (27; 14; 42), whereas classically activated microglia upregulate *P*2*X*4 and downregulate *P*2*Y* 12 (27). This motivated our hypothesis that P2X and P2Y co-expression in microglia subpopulations enable the cells to encode unique Ca^2+^ waveforms that prime migration versus inflammatory responses (Fig. 1).

**Figure 1:**
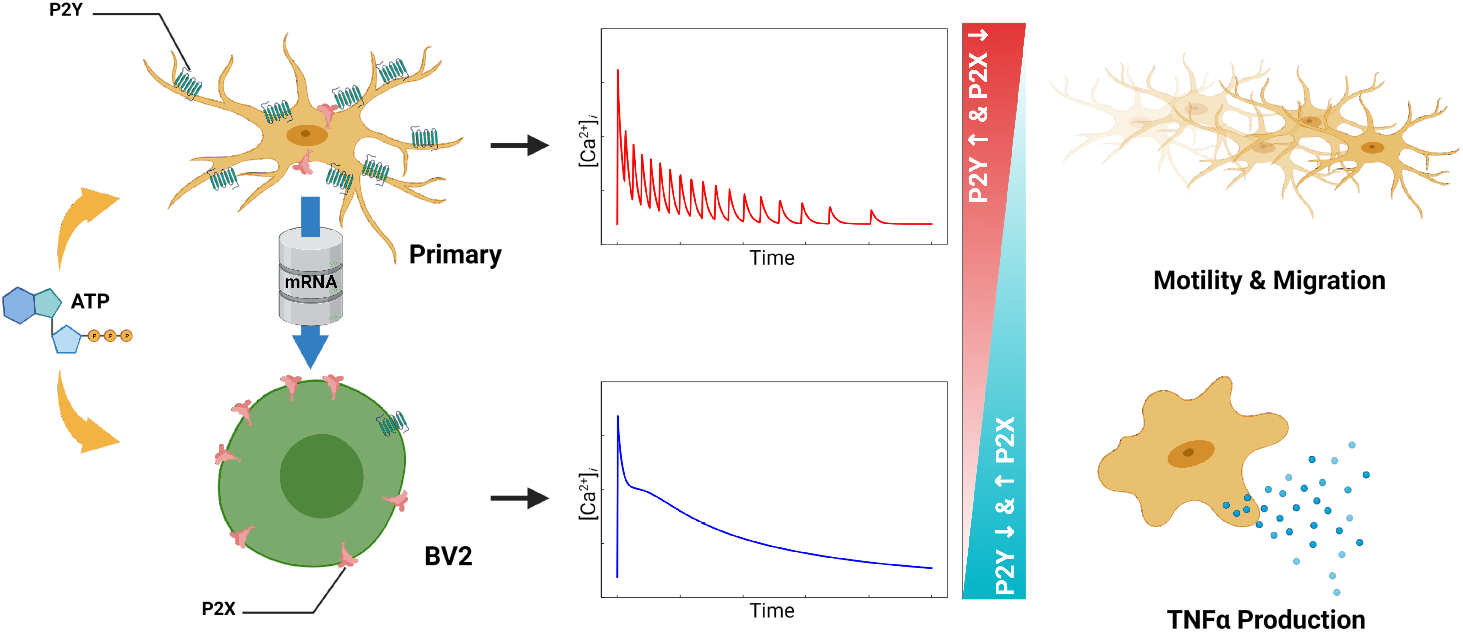
Table of Contents Graphic. Schematic of the computational study. ATP triggers the activation of P2Y and P2X receptors expressed in microglia (left column). The activated receptors generate intracellular Ca^2+^ signals (middle column) that contribute to microglial migration and cytokine production (right column). .

To investigate this hypothesis, we extended our model of *P*2*X*4/*P*2*X*7 activation in microglia (9) to include contributions from *P*2*Y*-class receptors. The extended model includes G-protein mediated Ca^2+^ signaling and the activation of pathways implicated in microglia migration. This approach complements prior computational studies of Ca^2+^ responses induced by *P*2*X* receptors (43; 9) and metabotropic receptors that promote intracellular Ca^2+^ release (16; 68; 67). With this contribution, we specifically examined how *P*2*X*- and *P*2*Y*-class purinoreceptors encode ATP-triggered Ca^2+^ waveforms in microglia, how these waveforms are modulated by enzymes like ectonucleotidases, and how these waveforms control migration and cytokine responses.

## 3 Materials and methods

### 3.1 The computational model

We extended the computational model of *P*2*X*4 and *P*2*X*7-mediated Ca^2+^ signals and TNF*α* production described in Chun *et al* (9) to include *P*2*Y*-dependent contributions to migration and cytokine responses (see Fig. 2). The original model consisted of differential-equation based descriptions of *P*2*X*4 and *P*2*X*7 gating (9) Ca^2+^ influx via *P*2*X* channels with the production of TNF*α*. The aforementioned model has been extended with the addition of pathways such as phosphoinositide 3-kinase (PI3K) activation and phosphorylation of Akt to provide a quantitative measure of microglial migration with respect to ATP (see Fig. 2) (59).

**Figure 2:**
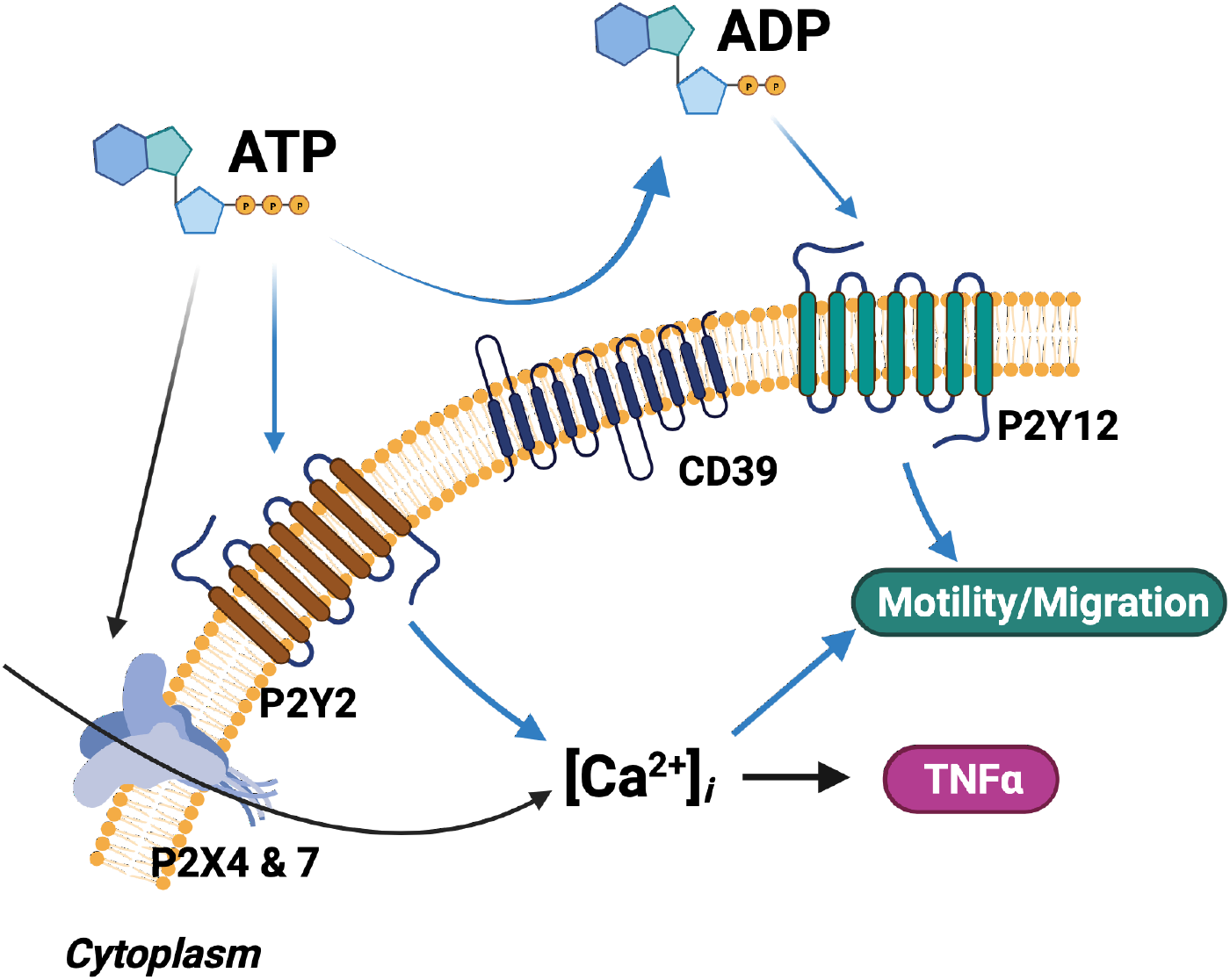
Schematic of the computational microglia model, which simulates the ATP-dependent activation of P2X- and P2Y-class receptors. The simulations also account for increases in intracellular Ca^2+^, cell migration, and TNF*α* cytokine production following P2 receptor activation. .

P2X4 and P2X7 models used in this study (see Fig. 9A) were implemented as described in Chun *et al* (9), which were originally derived from published models (85; 78; 43). Currents were related to intracellular Ca^2+^ by converting the inward current into the Ca^2+^ influx as described in our previous work (9).

For the metabotropic receptor contributions, we assumed that P2Y2 and P2Y6 activation promotes the disassembly of their *G_αq_* subunit, wherafter PLC is activated to promote the generation of Inositol trisphosphate (*IP*_3_) and Diacylglycerol (DAG) from PIP2 (42; 16) (see Fig. 9B). We described this process using a mathematical model introduced by Cutherbertson *et al* (16) for *IP*_3_-mediated Ca^2+^ oscillations in oocytes. The model includes an agonist receptor for *G_αq_* protein activation, *G_αq_*-dependent Phospholipase C (PLC) activation, PLC-dependent *IP*_3_ synthesis, and we also assume *P*2*Y* is activated by ATP, though it is likely that the activation occurs through the ADP product of ectonucleotidases (detailed in Sect. S.5.) DAG in turn indirectly inhibits G-protein dependent PLC activation by catalyzing PKC activity. The major challenge associated with validating contributions of specific *P*2*Y* receptors to experimentally measured Ca^2+^ transients is isolating the contribution of the specific receptor from other *P*2*Y* receptor contributions. We lump oscillatory transients into G-protein dynamics, PLC activity, and *IP*_3_-mediated Ca^2+^ release from the ER. Although a number of scientific communications have discussed the correlation between P2Y12/13 receptor activation and intracellular Ca^2+^ fluctuations via indirect pathways (40; 42), for model simplicity, we created a lumped *P*2*Y* model.*IP*_3_ molecules trigger the activation of *IP*_3_ receptors that induce ERCa^2+^ release. Concurrently, as the cytosolic Ca^2+^ rises following *IP*_3_ receptor opening, DAG and Ca^2+^ promote protein kinase C (PKC)-based inhibition of *G_αq_* and the P2Y receptors.

Ectonucleoside triphosphate diphosphohydrolase-1, also known as CD39, is expressed in the surface of microglial plasma-membrane and play an important role in microglial migration by balancing ATP and adenosine molecules (25). We used the model of ATP decomposition into ADP and AMP by CD39 introduced in the (47) (see Fig. 9C). We assume the agonist for P2X and P2Y-class that mediates Ca^2+^ fluctuations is ATP whereas P2Y12 is specifically stimulated by ADP.

Many models for cell migration have been reported in the literature. These includes models for actin polymerization (52; 64), multi-cellular migration (17), and tissue-level simulations of tumor growth (83). In microglia, migration is associated with pathways triggered by P2Y12 activation (see Fig. 9D). A G_*ai*_-containing GPCR, *P*2*Y* 12, ultimately promotes PI3K activation in response to ADP. Within the process of activating PI3K, the active form of PLC dephosphorylates PIP3, the product of which mediates the PI3K/Akt pathway (46). According to a series of work done by Ohsawa *et al* (58; 59), it is clear that the presence Ca^2+^ rise in inducing the chemotaxis of microglia is reflected as a sensitivity of PI3K activation to the intracellular Ca^2+^ concentration. In Ohsawa’s work(58), the inhibition of PI3K substantially reduces the phosphorylation of Akt, which not only results in the suppression of membrane ruffling and migration. In addition, we also introduce the involvement of calmodulin (CaM) that is an indirect component of chemotaxis process mediated via MLCK pathway, according to Yao *et al* (86).

For describing the dynamics of cytosolic Ca^2+^, we implemented and validated Ca^2+^ uptake and release from the ER by SERCA and *IP*_3_*R*, respectively. Basal Ca^2+^ levels are restored by the Sarcoplasmic/endoplasmic reticulum calcium ATPase (SERCA) pump and the sodium/calcium exchanger (NCX) as described in (9).

With respect to the ER Ca^2+^ load, we utilized ratiometric data from ATP-treated microglial cells presented by Ikeda *et al* (35) to estimate ER Ca^2+^ release via *IP*_3_*R* receptor (*IP*_3_*R*). The authors obtained a series of Ca^2+^ transients to infer the distinct contributions made by ionotropic and metabotropic receptors. The authors utilized two ATP concentrations (100 μM and 1 mM) to selectively activate a target receptor(s) (*P*2*X*4 vs. *P*2*X*7). In another experimental setup, *P*2*X*7R RNA*i* to silence *P*2*X*7 receptor that was used to isolate *P*2*X*7 contributions to the overall Ca^2+^ transients under constant ATP stimulation. For model validation, we converted the ratiometric data to concentration data by normalizing the cytosolic fluorescent intensity of resting microglia to 100 nM.

We neglect store operated calcium entry (*SOCE*) given that this mechanism of Ca^2+^ entry occurs well after Ca^2+^ currents mediated directly by 5-10 minutes after ATP stimulation(40; 35; 1; 5). Limitations of this simplification are discussed in Limitations (Sect. 6).

TNF*α* production via NFAT (see Fig. 9E) was implemented as described in Chun *et al* (9). Particular process we included in this work is that Ca^2+^-mediated CaM/calcineurin activation promotes the translocation of NFAT into nucleus (13), which mediates the transcription of TNF*α* mRNA (33).

#### 3.1.1 Numerical solution of the computational model

The resulting system of differential equations were numerically solved and optimized via Python (ver. 3.6) and Gotran (ver. 2020.2.0.), see (9) for details. As previously described (74; 9), the Generalized ordinary differential equations (ODE) (ordinary differential equation) Translator was utilized to implement the microglial model. The Gotran Python module was utilized to make use of our previously written Python-based routines for simulation and analysis. The SciPy function, ODEINT, that employs the LSODA algorithm for stiff ODEs (60) was used in the numerical integration of the microglia model. A time-step for the 10-min numerical integration was of 0.1 ms. These computations generate as output the time-dependent values of model ‘states’, such as intracellular Ca^2+^ or the open gates of the *P*2*X* channels. Model fitting was further tuned and refined by a genetic algorithm (reviewed in (71)) that iteratively improved assigned parameters, such as the rate of Ca^2+^ leak and *P*2*X*4/*P*2*X*7 conductance. Parameters for the model components are summarized in Sect. S.1. Based on these sets of parameters, our key model outputs were Ca^2+^ transients with respect to ATP exposure duration and concentration, as well subsequent changes in other states including PI3K and Akt for which experimental reference data were available. Experimentally-measured outputs, such as Ca^2+^ transient decay time and amplitude, were used to optimize the model parameters by minimizing the error between model predicted outputs and experiment.

All code written and simulation input files in support of this publication are publicly available at https://github.com/bending456/2021p2xp2y. Generated data are available upon request.

### 3.2 Experimental details

#### 3.2.1 Microglia Culture and imaging

For live cell calcium imaging, 96 well plates were used for BV2 cell culture. Cells were plated in a density of 5000 per wells 2 days prior to imaging. In the day of calcium imaging, the cells were incubated at 37°*C* with 5% *CO*_2_ with 1 *μg/mL* Fluo-4 AM (Thermo Fischer Scientific) in Leibovitz’s L-15 medium for 45 minutes. Excess Fluo-4 AM was washed by Leibovitz’s L-15 Medium and again incubated in Leibovitz’s L-15 Medium for 30 minutes. A custom-built, wide-field, epifluorescence microscope having a 10X objective with 488 nm laser was used for taking time lapse images. 20-minute time lapse movies were taken for different concentrations of ATP (ThermoFischer Scientific) with first 10 minute without adding ATP and rest after adding ATP.

For data in Fig. 7, Ca^2+^ measurements were taken every 0.5 mins throughout the experimental window. For each ATP dose, approximately 48 cells were selected to calculate the Ca^2+^ transients (as indicated by pixel intensity). For each cell, the ATP-induced Ca^2+^ transient was normalized with respect to the control stage, namely, the average pixel intensity of the control stage was subtracted from the pixel trace of the whole time course.

##### Detection of ATP-induced Ca transients

The recorded images via the aforementioned calcium imaging protocol were processed via custom python routines. The TIF file from experiment is processed as a 3D matrix with a specific shape defined by (T,M,N) where *T* denotes the time-index, *M × N* is the image dimensionality (in the unit of pixels). The pixel intensity (gray-scale) was stored in 16-bit color depth. As the first step, the cell detection protocol identifies cells from the image and provides its quantity. Specifically, the intensity at each pixel of the image was summed up along the time-index to get a total image with shape (*M × N*). The total image was then subject to a log transformation and normalization. The histogram of pixel intensities of the normalized image was plotted to help identify a custom thresholding value. Parameters used for acquisition of these signals are embedded in the source code provided with this project. After determining the thresholding value, the normalized total image was converted into a binary image with intensities greater than the thresholding value identified as cell bodies with the rest as background. The detected cells in the binary image satisfying the aforementioned condition were recognized and labeled. Using this information, the original TIF gray-scale image was utilized to record the change in the pixel intensity at each cell location, the average trace of which later represented the ATP-mediated Ca^2+^ transient of each cell.

We subsequently analyzed the Ca^2+^ transients before and after addition of different concentrations of ATP by taking time lapse images using widefield excitation. To normalize the data, we first calculated the average Ca^2+^ signal or fluorescent intensity in the control phase, the first 10 minutes without ATP. We then identified Ca^2+^ transients by dividing the entire time trace by the average fluorescent intensity. This was used to identify changes in the fluorescence signal.

##### Detection of BV2 motility

Using the bright-field data collected for ATP treated BV2 cells, we selected cells that exhibited displacements within the first 20 frames directly after ATP treatment, which corresponds to 600 seconds in total. To measure displacements, we manually selected reference points within a given BV2 cell process at the initial time point and its approximate position at the final (10th) frame, from which a vector was defined. The resolution of the brightfield data was approximately 1.6 μm/pixel, therefore the length in pixels of the displacement vector was converted into micrometers.

## 4 Results

### 4.1 Ca^2+^ waveform jointly is shaped by P2 receptors and ectonucleotidases

We developed a computer model of metabotropic (P2Y-class) and ionotropic (P2X-class) purinergic receptors to simulate Ca^2+^ signaling in microglia (Fig. 2). These Ca^2+^ signals are induced by extracellular ATP and its metabolites binding to P2 receptors (42; 56). We therefore expanded a published model of P2X receptor activation (9) to include *P*2*Y* in order to investigate how both purinoreceptor classes influence intracellular Ca^2+^ transients. We first implemented and validated a model for G-protein mediated *IP*_3_ generation and *IP*_3_ receptor-mediated ER Ca^2+^ release contributed by Cuthbertson *et al* (16) that we adapted to reflect P2Y-mediated Ca^2+^ waveforms (see Fig. 3). We then validated predicted Ca^2+^ waveforms generated by both P2X- and P2Y-class receptors against data collected in primary microglia by Ikeda *et al* (35).

**Figure 3:**
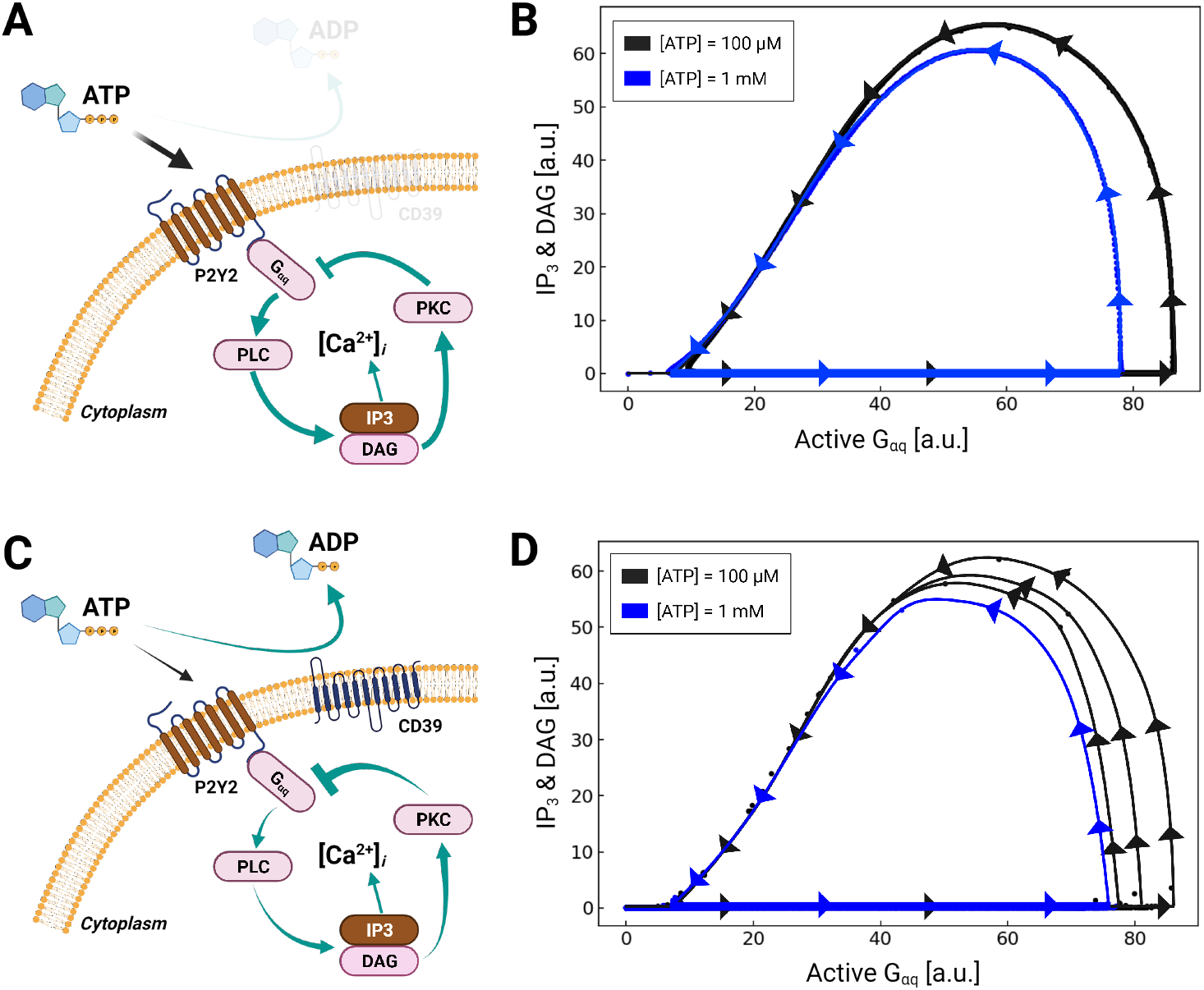
A) Schematic of P2Y2-dependent activation of *IP*_3_ production via *G_αq_*. B) Predictions of oscillatory *IP*_3_ and DAG production as a function of activated *G_αq_* in response to 100 μM (black) and 1 mM (blue) ATP. C) and D) are equivalent to A and B, except that the simulation includes the hydrolysis of ATP by the ectonucleotidase CD39. .

#### 4.1.1 *G_αq_* signaling via *P*2*Y*

*P*2*Y*-driven *IP*_3_ signaling begins with the activation of *G_αq_*-protein, which stimulates Phospholipase C (PLC) to produce Inositol trisphosphate (*IP*_3_) and Diacylglycerol (DAG) from *PIP*_2_ (Fig. 3 A). This is followed by ER Ca^2+^ release via *IP*_3_-stimulated *IP*_3_*R*s. Negative-feedback arises in this system as DAG produced by PLC activates Protein kinase C (PKC), which inhibits *G_αq_*. In Fig. 3B we demonstrate that the activation of *P*2*Y* by ATP and subsequently *G_αq_* results in periodic fluctuations in DAG and *IP*_3_ concentrations. This is evident as stationary cycles in Fig. 4A, where increases in active *G_αq_* were accompanied by increases in *IP*_3_ and DAG; these increases continued until active *G_αq_* was nearly saturated at 80 [a.u], whereafter *IP*_3_ and DAG rapidly decayed to zero as active *G_αq_* was depleted. Larger oscillations in *G_αq_* activation and *IP*_3_ production were evident with 1 mM ATP relative to 1 μM.

**Figure 4:**
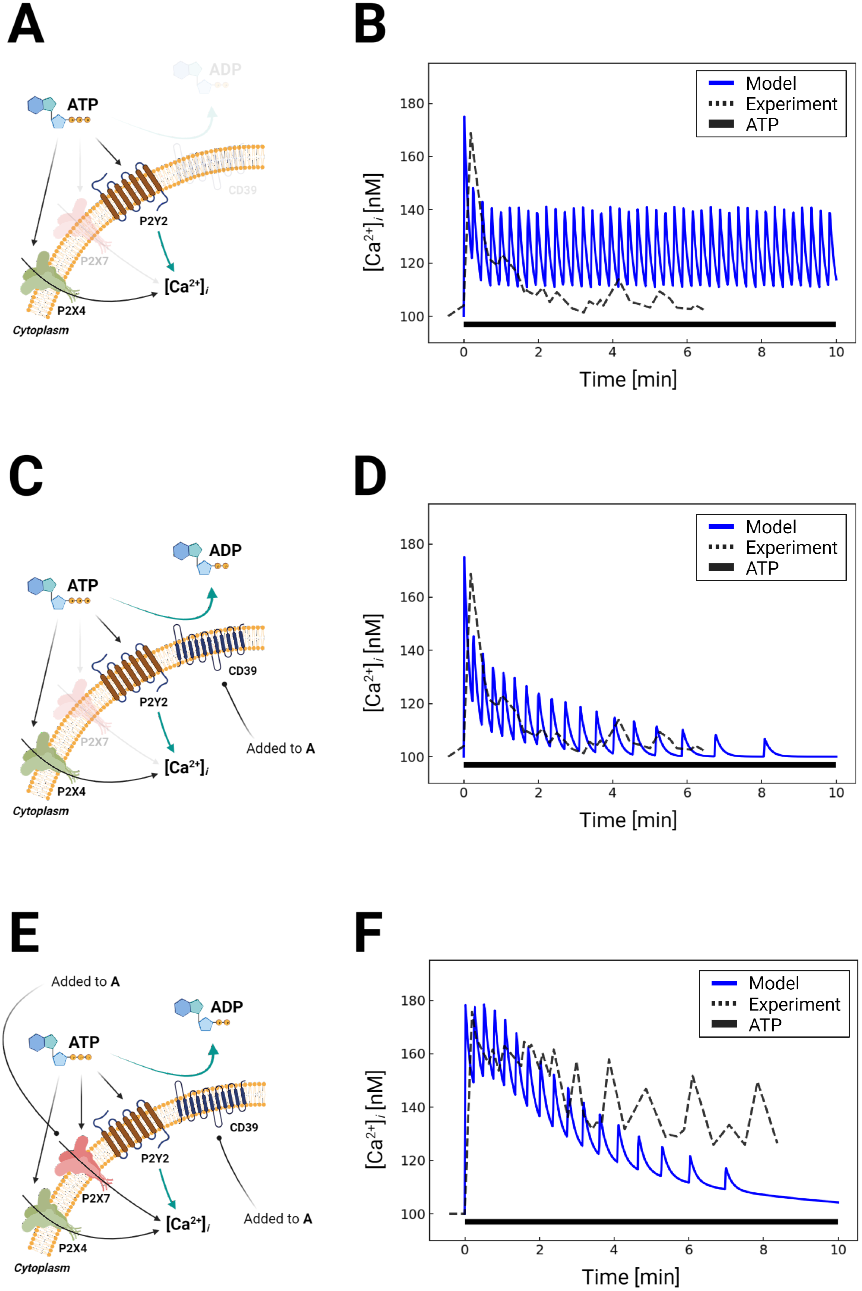
A) Schematic for Ca^2+^ waveforms generated by P2X4 and P2Y2 in response to 1mM ATP for 5 minutes. P2X7 contributions were blocked in this simulation (P2X7 KO). Schematic C) is analogous to A), but includes CD39 nucleotidase degradation of ATP (P2X7 KO). E) Schematic for Ca^2+^ waveforms generated in a control system comprising P2X4, P2X7 and P2Y2 and includes CD39 nucleotidase activity (WT). B, D, and F) Comparison of predicted (blue) and experimentally-measured (35) (dashed) Ca^2+^ transients, corresponding to the schematic A, C, and E, respectively. Results excluding CD39 are shown in Fig. S6.

Stable *IP*_3_ oscillations are reported in many cells types and are a prototypical example of negative-feedback circuits (82; 41). Interestingly, microglia exhibit both oscillatory and aperiodic Ca^2+^ transients, which suggests that the underlying *IP*_3_ synthesis is not exclusively periodic. Oscillations in feedback biochemical circuits are determined by the kinetics of the underlying enzymes, therefore non-oscillatory *IP*_3_ signals are theoretically possible and would manifest as single-peak Ca^2+^ ‘waveforms’ (35). We demonstrate in Fig. S3) how variations in the parameters underlying P2Y-frequency dependent *G_αq_* activation can yield stable oscillations versus aperiodic behavior. Namely, by reducing the input parameter *kg_p_*_2*y*_ that controls the rate of *G_αq_* activation, the system reverts to nonoscillatory behavior. Similar effects can be shown by varying other parameters describing the *G_αq_* negative-feedback circuit, which suggests that the activity of proteins comprising the *G_αq_*/*IP*_3_ signaling pathway determine whether *IP*_3_ and thereby Ca^2+^ are oscillatory versus non-oscillatory.

*IP*_3_ invokes intracellular Ca^2+^ release from the endoplasmic reticulum via *IP*_3_receptors. We therefore fit the model’s predicted *IP*_3_-induced Ca^2+^ transients to reproduce experimental data collected by Ikeda *et al* (35). Namely, we fit the initial peak amplitude ([Ca^2+^]_*i*_ = 180 nM) to match the Ikeda *et al* data in MG5 microglial cells treated with 1 mM ATP (35), for which *P*2*X*7 was knocked-out to isolate P2Y contributions. We reflected this condition in our overall Ca^2+^ signaling model by disabling the *P*2*X*-mediated currents. In Fig. 4b, we compare our predictions of *IP*_3_-mediated Ca^2+^ release following a 10-minute 1 mM ATP treatment (blue) relative to the experimentally-measured transients from Ikeda *et al* (dashed black). After this initial peak, our model predicts an oscillatory Ca^2+^ waveform that is complemented by decreases in ER Ca^2+^ owing to *IP*_3_ receptor activation (Fig. S2). While the predicted Ca^2+^ waveform resembles the oscillations observed by Ikeda *et al*, the waveform did not exhibit decay with time, in contrast with experiment.

While a variety of mechanisms could be attributed to this discrepancy, such as the desensitization of *P*2*Y* receptors to ATP (38), we speculated that the availability of ATP for triggering *P*2*Y* was the prominent source of error. Normally, extracellular ATP is rapidly degraded into adenosine diphosphate (ADP) and adenosine monophosphate (AMP) by ectonucleotidases (47). In microglia, NTPDase1 (CD39) is the primary ectonucleotidase isoform responsible for ATP degradation (25). To represent this contribution, we implemented a mathematical model from (47) to simulate ectonucleotidase-catalyzed hydrolysis of ATP into ADP and AMP. Our implementation is validated against experimental data in Fig. S4, for which nucleotide concentrations were measured in COS-7 cells over a one-hour time interval (47). Importantly, these data demonstrate that the ATP pool was depleted within minutes; this depletion was accompanied by a transient ADP pool that was maximal at t=4 min and subsequently decayed to zero. After including ectonucleotidase contributions in our microglia model, the predicted cytosolic Ca^2+^ transients decayed in a manner consisted with experiment (Fig. 4B) without additional fitting. Interestingly, we found that the frequency of predicted oscillations decreased with time, which we attributed to the dose-dependent decrease in *G_αq_* activated by the P2Y receptors (Fig. S3). Hence, our simulations provide strong evidence that ectonucleotidases, and specifically CD39, play a prominent role in shaping the Ca^2+^ waveform by controlling the nucleotide pool available to purinoreceptors.

After validating our model of P2Y-induced Ca^2+^ dynamics, we restored P2X receptor contributions and compared model predictions against analogous experiments in Fig. 4. We again simulated the system subjected to 1 mM ATP for ten minutes with and without ectonucleotidase activity (shown in Fig. 4 and Fig. S6). In contrast to the *P*2*X*7 knock-out data, we observed a modestly higher peak Ca^2+^ transient amplitude that was followed by a prolonged plateau as would be expected from *P*2*X*7 currents (20). Predicted Ca^2+^ oscillations decayed toward resting Ca^2+^ levels after approximately ten minutes, albeit at a faster rate than observed experimentally. We report similar findings upon 100 uM ATP treatment in Fig. S5, which favored *P*2*X*4 activation. Altogether, our data suggest that a diverse ensemble of Ca^2+^ waveforms are invoked by controlling P2 receptor activation and nucleotide availability.

### 4.2 P2 mediated Ca^2+^ waveforms contribute to migration

We next examined how *P*2*Y* activation and *P*2*Y*-mediated Ca^2+^ waveforms control cell migration and motility (see Fig. 5). *P*2*Y* 12 activation is essential for chemotactic migration and rapid motility responses in ATP-stimulated microglia (58; 59; 36; 42). *P*2*Y* 12 receptors primarily activate the G-protein, G_*ai*_, and thereby promote Akt phosphorylation via PI3K, which in turn activates mechanisms underlying cell migration in microglia (59).

**Figure 5:**
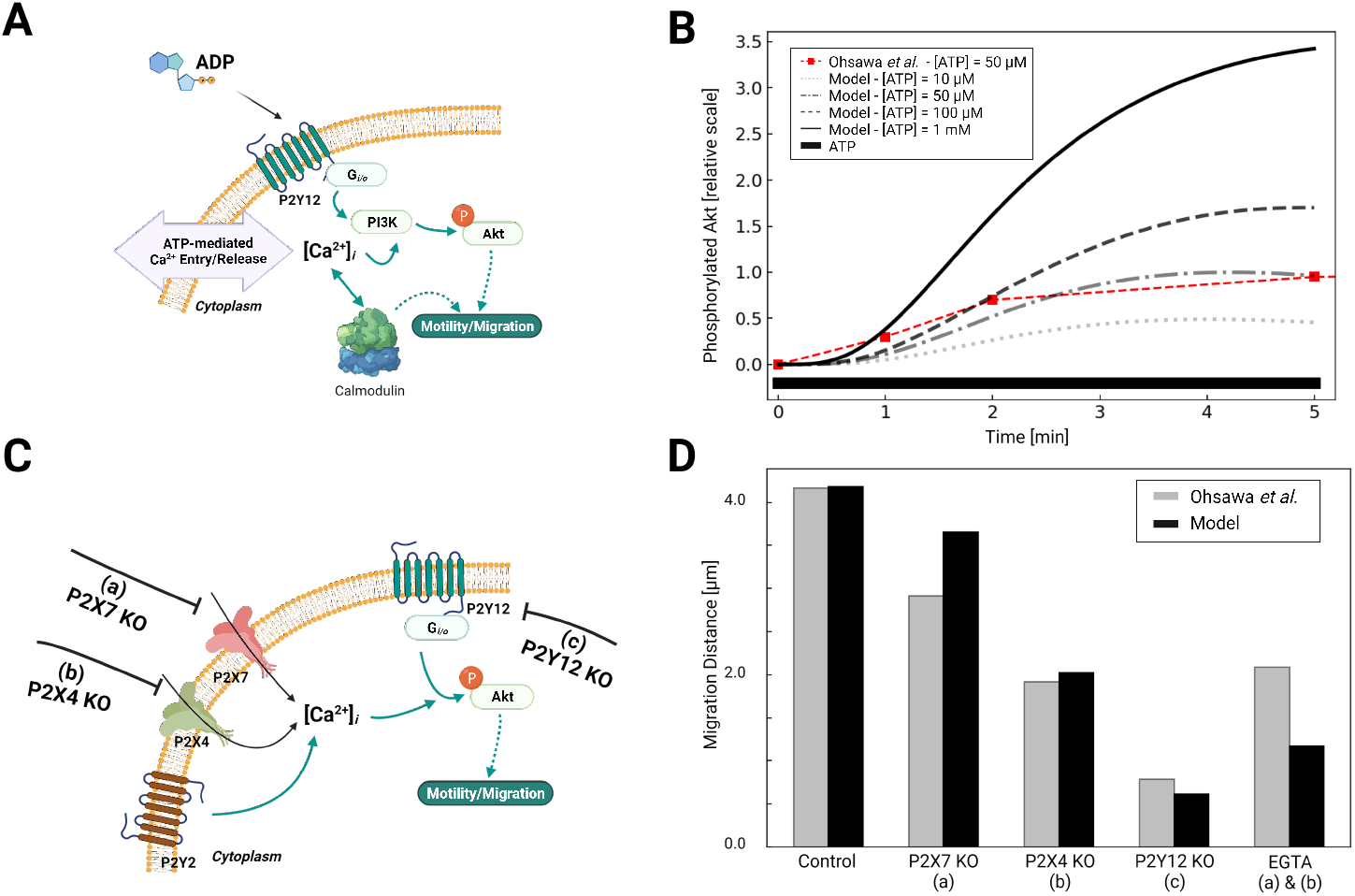
A) Schematic of Akt phosphorylation via P2Y12- and P2Y-mediated Ca^2+^ signaling pathways in response to ATP. B) Predicted *p*Akt expression as a function of time in response to 50 uM to 1 mM ATP applied for 5 minutes. Experimental data for 50 uM ATP from Ohsawa *et al* are presented in red. C) Schematic of P2Y12- and Ca-mediated migration in response to ATP, assuming control, P2X7 knockout (KO) (a), P2X4 KO (b) and P2Y12 KO (c) conditions. D) Predicted migration distances (black) versus experimental measurements by Ohsawa *et al* under control and a-c conditions. .

We first fitted our *p*Akt responses under 50 μM ATP treatment to reproduce trends observed by Ohsawa *et al*. For this process, we utilized the time-dependent phosphorylation of Akt following the activation of PI3K and the data with various P2 KO conditions to tune the sensitivity of PI3K/*p*Akt pathways to the intercellular Ca^2+^ fluctuations. The resulting fit is shown in Fig. 5A, which indicates maximal *p*Akt levels were obtained at about 4 minutes and were in close agreement with experiment. *P*2*Y* 12 activation peaked at this time given that its primary agonist, ADP, is maximal at 4 minutes due to ectonucleotidase activity (see Fig. S4). We show that Akt phosphorylation increased with increasing ATP concentration, as 100 μM and 1 mM ATP treatments resulted in 50 and 300% increases in pAkt relative to the 50 μM treatment.

Curiously, this process has been shown to depend on extracellular Ca^2+^, as inhibition of *P*2*X* receptors (PPADS and TNP-ATP for *P*2*X*7 and *P*2*X*4, respectively) and chelation of extracellular Ca^2+^ by EGTA all diminish both *p*Akt levels and migration (58; 59; 36). We therefore augmented our computational model to include Ca^2+^-dependent PI3K/*p*Akt activation, as a step toward investigating the extents to which purinoreceptor-encoded Ca^2+^ waveforms influence migration (18). To determine the Ca^2+^ dependence of PI3K activation, we referred to data from Ohsawa *et al* (58) that reported Akt phosphorylation following *P*2*Y* 12, *P*2*X*7 or *P*2*X*4 knockout in neonatal Wistar rat microglia. Under *P*2*Y* 12 knock-out, they observed a 90% reduction in *p*Akt upon treatment of 50 μM ATP for 5 mins relative to WT. We attributed the remaining 10% of the *p*Akt phosphorylation to Ca^2+^ influx from the P2X receptors. This was motivated by our observations that 1) *P*2*X*4 in particular generated prominent Ca^2+^ transients with micromolar ATP treatments (59) and 2) that EGTA treatment nullified *P*2*X*-mediated Ca^2+^ transients and significantly reduces PI3K activation (59). Indeed, we predict in Fig. S8C that EGTA treatment reduced *p*Akt by 75%, which implicated extracellular Ca^2+^ as a significant contribution to microglia migration.

We next examined the effects of *P*2*X*4 and *P*2*X*7 inhibition on PI3K activation, and subsequently, *p*Akt, as both conduct extracellular Ca^2+^. Without additional refitting our model predicted 15% and 40% reductions in *p*Akt levels solely from *P*2*X*4 and *P*2*X*7 inhibition, respectively, in close agreement with Ohsawa (see Fig. 5B), These significant reductions upon nullifying *P*2*X* contributions therefore implicate ionotropic receptors in phosphorylating Akt. We additionally verified that *P*2*Y* 12 knock-out all but eliminates Akt phosphorylation. Lastly, we predict that blunted ectonucleotidase hydrolase function enhances *p*Akt phosphorylation, which suggests that prolonged Ca^2+^ waveforms further promote Akt activation (58) (shown in Fig. S8B).

#### 4.2.1 Microglial migration

*P*2*Y* 12, PI3K, Akt and extracellular Ca^2+^ are necessary for migration (58; 59). This is in part supported by data from Ohsawa *et al* demonstrating microglia with inhibited PI3K exhibit in negligible migration when treated with ATP (58). We therefore assumed microglia migration rates were proportional to *p*Akt levels in accordance with data from Ohsawa (58). Those data suggested that ATP-treated microglia migrate distances of approximately 48 micron following one hour of 50 μM treatment. We interpolated these data to a distance 4 μm following five minutes of stimulation. The *p*Akt-dependent migration rate was then fit to yield the this short migration distance following integration of the *p*Akt levels over five minutes. We tested our fitted model by reporting migration distances upon inhibition of *P*2*X*4, *P*2*X*7, and *P*2*Y* 12 (see Fig. 5C). Without additional fitting, our simulated data nearly reproduced the 75% reduction in migration following *P*2*Y* 12 knock-out that was reported by Ohsawa *et al*. Intermediate reductions in migration distances following *P*2*X*7 and *P*2*X*4 knockout were also comparable to data from Ohsawa *et al* (Fig. 5D). We further demonstrate that reducing ectonucleotidase activity enhanced migration, as was already observed for our predicted *p*Akt levels and Ca^2+^ waveform durations (shown in Fig. S9). Importantly, these simulated data confirm that *P*2*Y* 12 KO dramatically reduced, but did not entirely eliminate migration; together with the reductions in migration following *P*2*X* knock-outs, these data implicate the significant role of extracellular Ca^2+^ in mediating migration.

### 4.3 Ca^2+^ waveforms and their impact on migration versus cytokine responses

Our data thus far indicate that microglia have robust Ca^2+^ responses to P2 receptor activation that promote migration. We previously showed in (9) that cytokine synthesis and release in microglia was driven by intracellular Ca^2+^ signaling. This raised the question as to how ATP-induced Ca^2+^ waveforms determined migration versus inflammatory cytokine responses in microglia. To answer this question, we predicted migration distances as a function of Ca^2+^ waveform amplitude and oscillation frequency when subject to 200 uM ATP for 5 minutes in Fig. 6. The amplitudes and frequencies were controlled by modulating the *IP*_3_ pathway parameter *kg_p_*_2*y*_. As a measure of cytokine responses, we report predicted TNF*α* released levels, using our validated model from (9). For reference, the dashed box indicates baseline migration and TNF*α* responses for the model when subject to 200 μM ATP. Per (33) *et al*, secreted TNF*α* was undetectable under 1 hr and reached a maximum concentration (425 pg/10^6^ cells) for 3 mM ATP after 6 hours. Our model predictions indicate that both migration and TNF*α* mRNA levels increase with increasing Ca^2+^ waveform frequency and amplitude (Fig. 6). Although TNF*α* responses were predicted to increase at a greater rate than migration for increasing Ca^2+^ waveform frequency and amplitude, we anticipate produced TNF*α* would nonetheless be undetectable at 5 minutes. This is based on observations by Hide *et al* that minimal TNF*α* (10 pg/10^6^ cells) was measured at 1 hr, even with ATP treatments exceeding 1 mM (3 mM). Importantly, our model demonstrates that 1) both migration and TNF*α* production were positively correlated with the frequency and amplitude of the Ca^2+^ waveform and 2) migration was induced insignificant TNF*α* responses when the Ca^2+^ waveforms were of short duration, low amplitude and low frequency.

**Figure 6:**
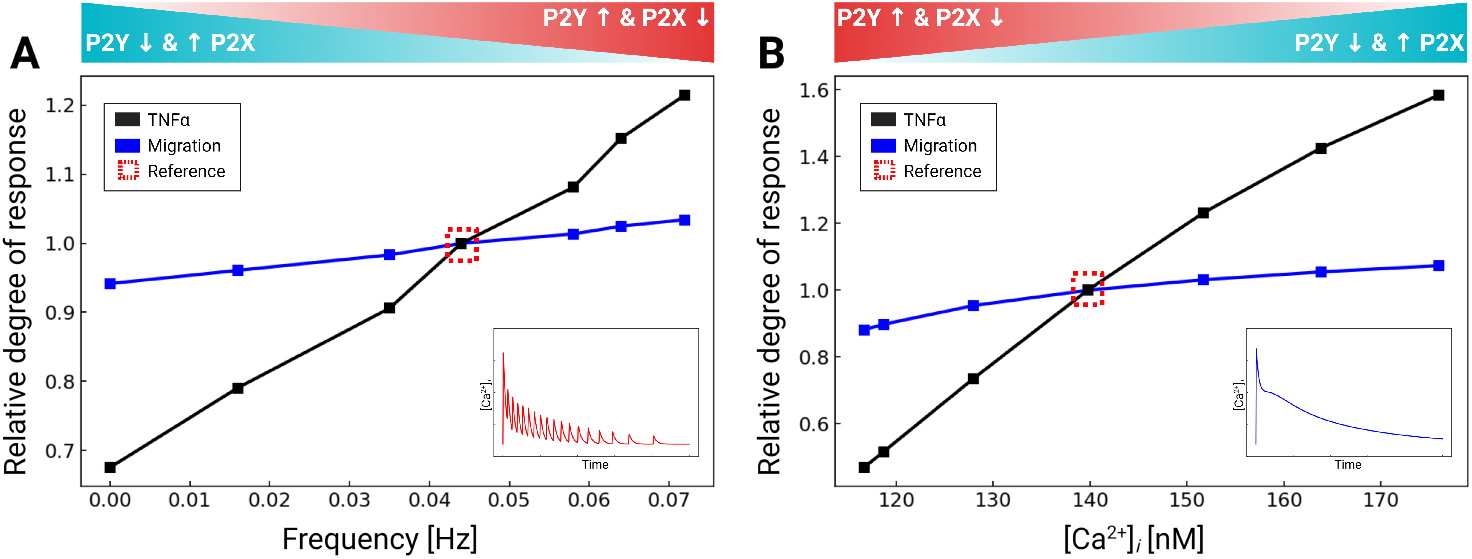
Predicted TNF*α* production (blue) and migration (black), normalized to control conditions (dashed box) in response to variations of the intra-cellular Ca^2+^ waveform frequency A) and amplitude B). All calculations were performed using 200 μM ATP, and the data were collected at 5-min. .

### 4.4 ATP-dependent responses in different microglia phenotypes

Microglia and their immortalized cell lines assume diverse phenotypes that are often characterized by differences in purinoreceptor expression (30; 5; 15). We therefore tested how such changes in purinoreceptor expression influenced ATP-triggered Ca^2+^ waveforms and Ca^2+^-dependent functions. Although mRNA expression levels do not necessarily directly correlate with protein expression (21), as a first approximation we proportionately rescaled the purinergic receptor responses in our model according to the relative change in mRNA expression (see Table 1). We adjusted P2 contributions in our model in accordance with mRNA data sets published for P2X4, P2X7, P2Y2 and P2Y12 (see Fig. 7) in primary relative to BV2 microglial cells (30). Those data reflect 5-fold and 2-fold reductions in *P*2*X*4 and *P*2*X*7 mRNA relative to primary cells, no change in P2Y2, and a near complete elimination of *P*2*Y* 12 mRNA.

**Table 1:** Relative expression of P2 receptors in acutely isolated, primary cultured microglia, and BV2 cells in relative scale. Reported receptor mRNA expression is normalized to the expression level (mRNA count) found in primary cultured microglia and are incorporated in our model as scaling factors for receptor concentration (*ρ_P_*_2*X*4_, *ρ_P_*_2*X*7_, *ρ_P_*_2*Y c*_, and *ρ_P_*_2*Y* 12_). *The mRNA expression was acquired from the comparison between cultured mouse microglia and BV2 cells(30).

| Receptors | mRNA Expression |  |
| --- | --- | --- |
|  | Primary | BV2* |
| *P2X4* | 1× | 0.18× |
| *P2X7* | 1× | 0.65× |
| P2Y2 | 1× | 1.02× |
| P2Y12 | 1× | 0.030× |

**Figure 7:**
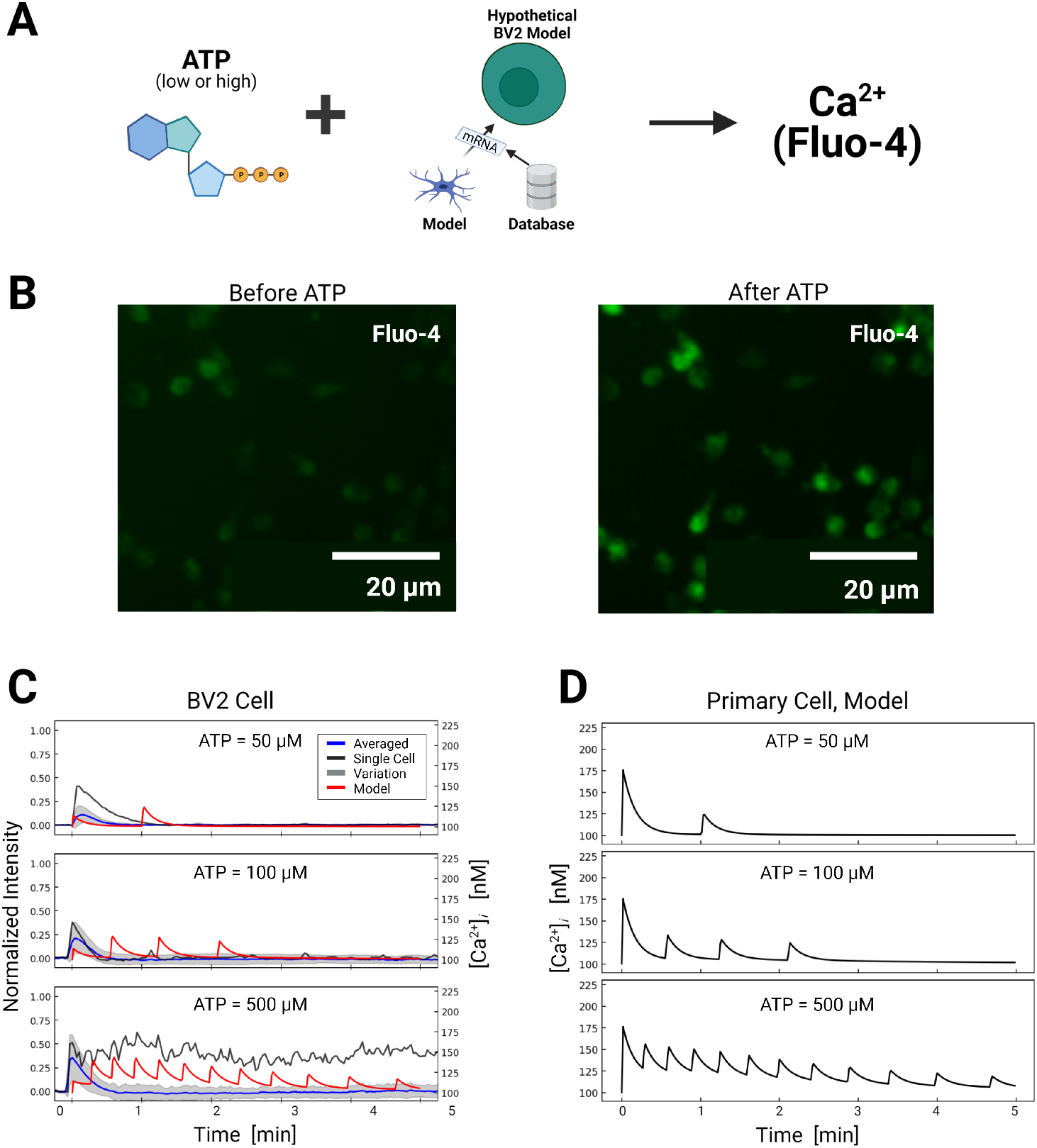
A) Schematic of ATP-induced Ca^2+^ transients in model BV2 cells assuming P2 expression levels inferred from BV2 cell mRNA (30). B) Ca^2+^ transients measured in BV2 microglia just before and after ATP treatment. C) Experimentally measured (blue) and predicted (red) Ca^2+^ transients from BV2 microglia. The shaded area represents the variation in signals measured in all cells. Oscillatory calcium signals were observed and a representative example of which is reported for each ATP dose (black). Transients recorded in each cell are reported in Fig. S12. D) Predicted transients in primary microglia for comparison. .

#### 4.4.1 BV2 Ca^2+^ transients

Based on the adjusted P2 responses, and without any additional fitting, we predicted Ca^2+^ transients in response to 50-500 μM ATP treatment applied for 10 mins (Fig. 7). The model demonstrated that the reduced *P*2*X* expression implied for BV2 cells resulted in moderately smaller Ca^2+^ transients relative to primary cells. The largest reductions were predicted at 50 and 100 uM, which was consistent with the preferential activation of *P*2*X*4 with micromolar ATP and the channel’s five-fold reduction in mRNA versus primary microglia. In contrast, modest reductions of 15% in Ca^2+^ transients after 20 seconds were predicted at 500 μM, which was inline with the *P*2*X*7 channel’s 35% reduction in *P*2*X*7 mRNA expression. Oscillations stemming from P2Y2 activation were predicted in the BV2 line and commensurate with those from primary cells. Importantly, the model confirms the intuitive result that BV2 Ca^2+^ transient amplitudes were reduced in a manner consistent with 5- and 2-fold reductions in *P*2*X*4 and *P*2*X*7 mRNA.

To validate these model predictions, we measured Ca^2+^ transients in cultured BV-2 cells. Because we did not have calibrated BV2 Ca^2+^ data, we assumed the peak Ca^2+^ amplitudes at 50 μM were approximately 112 nM in amplitude to be consistent with the 82% reduction *P*2*X*4 mRNA. We report in Fig. 7 the average Ca^2+^ transients (black) for the BV cells, while the cell-to-cell variance in represented by a gray shaded region. We found that the initial phase (<2min) of the predicted Ca^2+^ transients at 100 and 500 uM were in strong agreement with the experimental data. However, the cell-to-cell average did not exhibit fluctuations predicted by our model. Given that the mRNA data suggested similar P2Y2 expression in BV2 cells compared to primary cells, the model disagreement suggests that either ATP availability (as controlled by ectonucleotidase activity) or components of the *IP*_3_ Ca^2+^ signaling pathway may be attenuated. We were unable to evaluate this assumption as mRNA data were not available for the corresponding proteins. Interestingly, we included for reference experimentally-measured Ca^2+^ ‘outliers’ (blue) that strongly diverged from the population average and exhibited weak Ca^2+^ oscillations, which suggested that a subset of the cells had intact oscillatory *IP*_3_ signaling. Overall, it was evident from our model predictions that the expression differences in P2X channel mRNA were sufficient to reproduce the initial phase of the experimentally-measured Ca^2+^ transients and capture Ca^2+^ oscillations evident in a subset of BV2 cells. However, additional measurements of protein mRNA or expression levels of P2Y2 or downstream targets would ultimately be necessary to align the model predictions with experimental observations.

#### 4.4.2 BV2 migration

We last predicted BV2 migration upon 5 min ATP treatment intervals, based on the assumptions of reduced *P*2*X* and *P*2*Y* 12 expression (see Fig. 8). In accordance with the reduced *P*2*Y* 12 mRNA measured in BV2 cells, across all ATP concentrations we predicted a nearly 70% reduction in migration relative to primary cells. This reduction was consistent with the *P*2*Y* 12 knock-out data reported by Ohsawa *et al* that demonstrated reduced, but not entirely eliminated, migration in primary microglia. The predicted distances monotonically increased with higher concentrations of applied ATP, which was consistent with the Ca^2+^ dependency in migration exemplified in Fig. 6.

**Figure 8:**
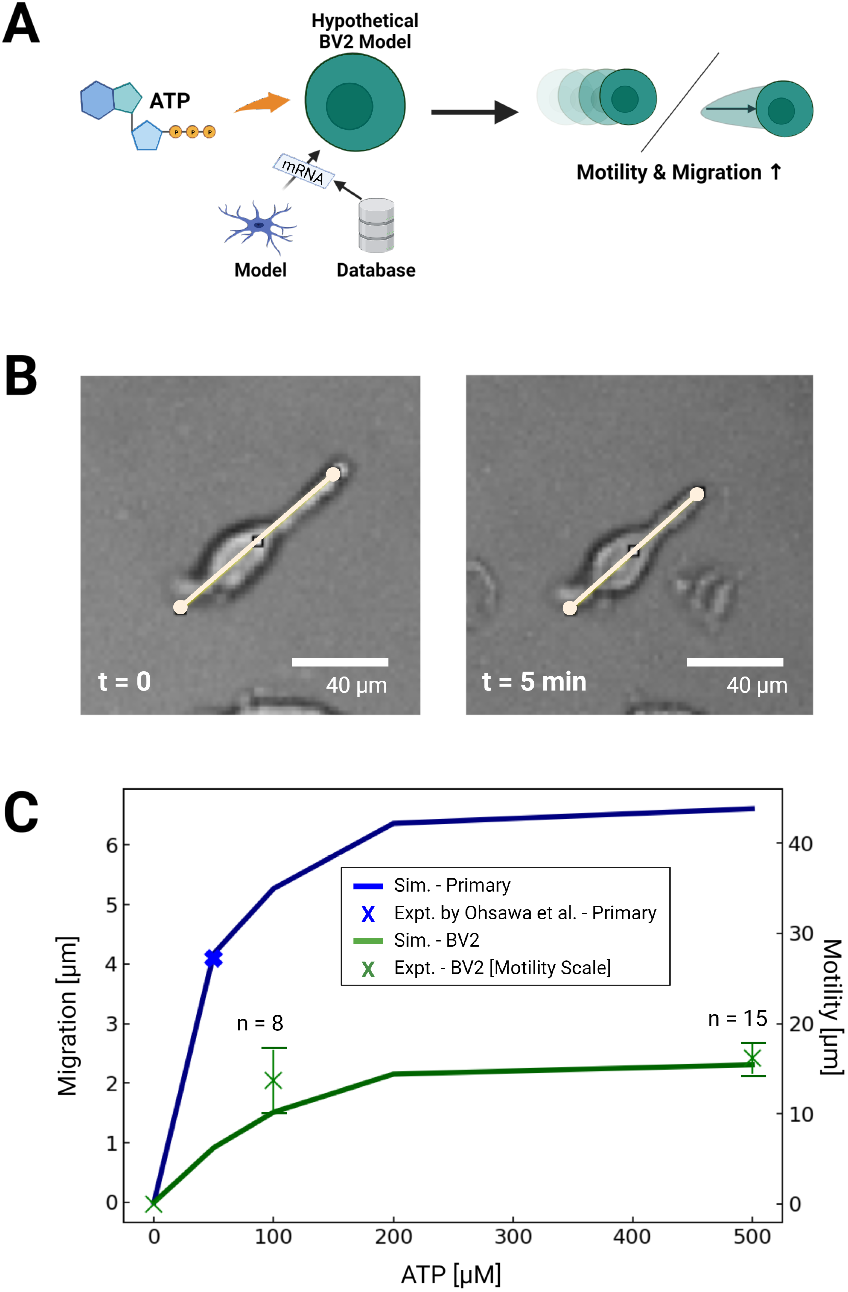
A) Schematic of predicted migration or motility in primary microglial cells or cells with P2 expression estimated from BV2 cell mRNA (30). B) Bright field image demonstrating ATP-dependent motility of microglial cell membrane (100 uM ATP at 5 minutes) C) Predicted (lines) migration or motility distance following 5 minute ATP treatment (0-500 uM) in primary microglia (blue) and BV2 cells (green). Experimental data in primary cells from Ohsawa *et al* and measured in BV2 cells are shown by blue and green X’s, respectively, where the bars represent standard error. see notes below

To validate these predictions, we examined subsets of BV2 cells that exhibited linear extensions of their membrane akin to the cellular processes evident in branched microglia (shown in Fig. 8B). We found that these cells rapidly contracted upon ATP treatment, as exemplified in Fig. 8, but otherwise we did not observe appreciable directed migration over the data collection interval. We there defined this motility as the displacement of plasma membrane following ATP treatment. To assess ATP-dose dependencies for these responses, we measured the displacement of these extensions along manually-defined vectors. We report in Fig. 8C that maximum displacement distance of 1.6 × 10^1^ μm were evident in response to 500 μm ATP,. Further, the dose-dependent displacement rates were consistent with the migration distances we predicted for BV2 cells in Fig. 8. Although these displacements were only reflected in a minority of the imaged cells, we found that morphological changes, surface ruffling and minor displacements of the cell somas were evident in a larger number of cells by visual inspection. These effects were more prominent for ATP-treated cells than those treated with saline (ATP = 0 uM). Overall, our quantitative and qualitative assessments of BV2 motility were consistent with data from Zhang *et al* (91) that demonstrated BV2 cells exhibit a two-fold migration ‘index’ after 24 hours when subject to ATP relative to to control conditions.

## 5 Discussion

### 5.1 Findings of this paper

In this study we used computational modeling to investigate how P2X and P2Y receptors collectively regulate Ca^2+^ and migration in microglia. These simulations indicate that *P*2*X* and *P*2*Y* encode Ca^2+^ signal waveforms that can selectively promote migration versus TNF*α* responses to ATP. This investigation necessitated extending a computational model of ionotropic *P*2*X* receptor activation (9) to incorporate contributions from metabotropic *P*2*Y* receptor activation. A schematic of the resulting model is shown in Fig. 9. With this model, we examined how the Ca^2+^ waveform from *P*2*Y* receptors differ from those generated via ionotropic means, how Ca^2+^ transient waveforms influence migration and TNFa responses, how those processes are shaped by the relative activity of *P*2*X* and *P*2*Y* receptors, as well as nucleotidases, and how purinergic receptor mRNA data could be used to extrapolate the model to other microglial cell phenotypes.

**Figure 9:**
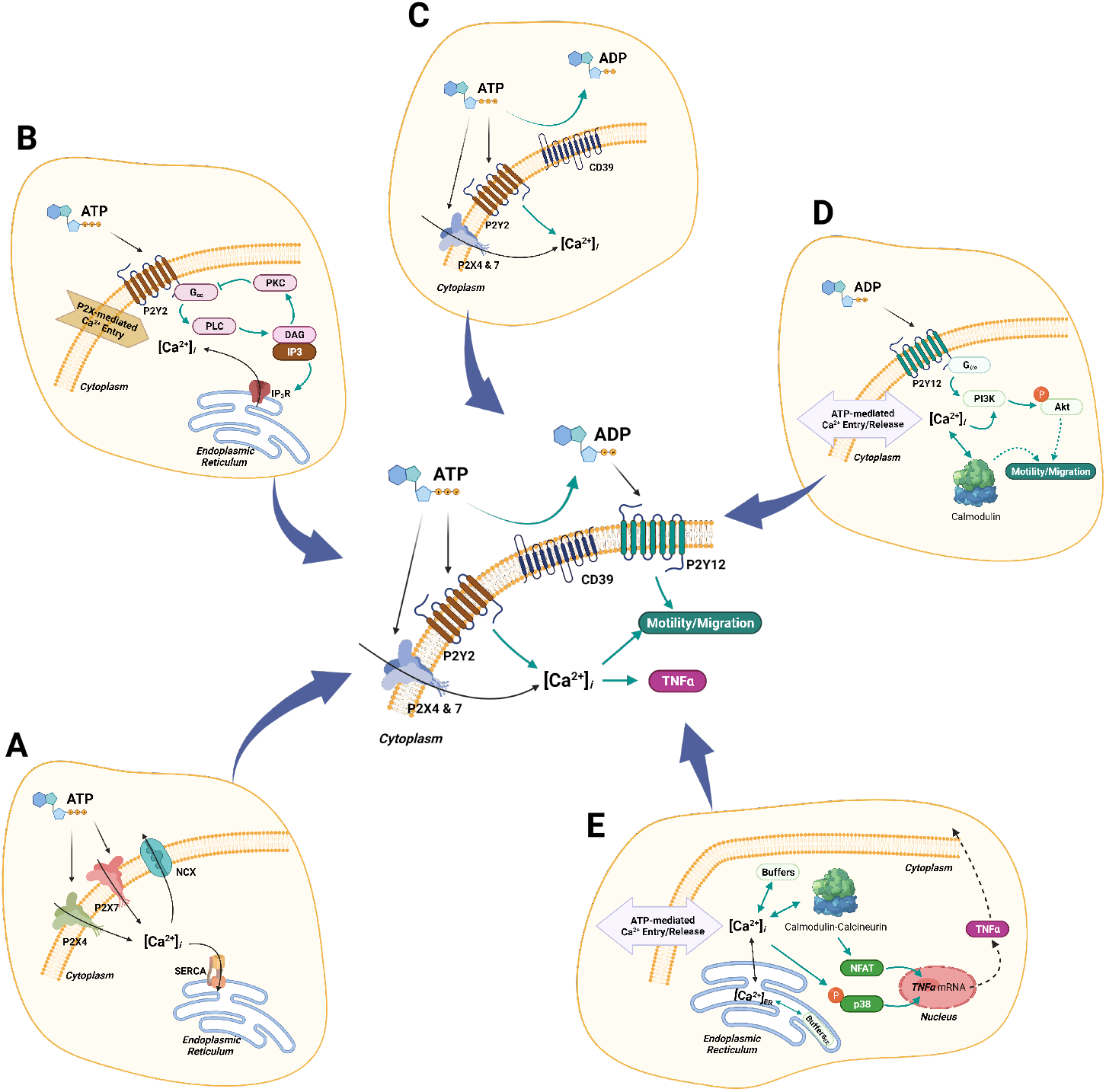
Complete schematic of the computational model. A) P2X4/7-mediated Ca^2+^ entry. B) P2Y2-mediated Ca^2+^ transients. C) Hydrolysis of ATP by ENTs (CD39). D) P2Y12-mediated cell migration/motility. E) Ca^2+^-dependent downstream cascades associated with TNF*α* production. .

### 5.2 Purinergic receptors and ectonucleotidases collectively control microglial Ca^2+^ waveforms

Our first goal was to determine the extent to which highly-expressed P2 receptors in microglia shaped the waveform of intracellular Ca^2+^ signals in response to ATP. Indeed, our model indicated that metabotropic P2Y receptors contribute significantly to ATP-induced Ca^2+^ signals in microglia. Since *in vivo* microglia typically express high levels of *P*2*Y* receptors (14), this further suggests that ER Ca^2+^ release may play a more significant role in tissue microglia than would be observed in *ex vivo* cultured microglia that are more commonly studied. Further, unlike the P2X receptors that generally present high-amplitude, single-peak Ca^2+^ waveforms, we show that P2Y receptors can adopt oscillatory or transient, single-peak, Ca^2+^ fluctuations (shown in Fig. S3). Given that sustained increases in basal intracellular Ca^2+^ levels are associated with pathological states (81; 34) and spontaneous oscillations are typical of homeostatic cells (42; 34), the balance of P2X versus P2Y contributions likely helps determine or at least indicate the cell phenotype (Fig. 10). Additionally, the ability for P2Y to encode diverse oscillatory and nonoscillatory signals could serve as a mechanism for controlling Ca^2+^-dependent functions in microglia. Our speculation is consistent with findings in other Eukaryotic cells that the dynamic profiles of Ca^2+^ waveforms tune cellular outcomes (69). As examples, oscillatory Ca^2+^ waves in oocytes are observed during in fertilization, while the timing of Ca^2+^ pulses in cardiac myocytes can selectively activate rapid CaMKII-versus slow NFAT-mediated gene responses (89).

**Figure 10:**
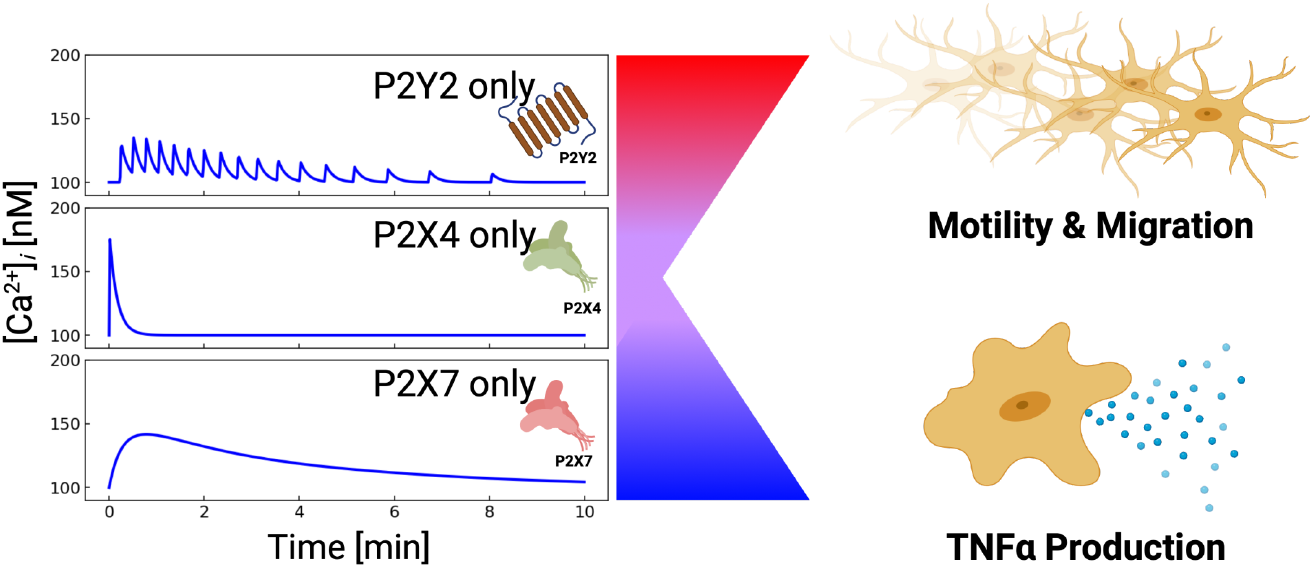
P2s can be combined to control different cellular outcomes. *P*2*X*4-induced Ca^2+^ entry manifests a sharp but brief rise in cytosolic Ca^2+^. *P*2*X*7-mediated Ca^2+^ entry results in blunt and prolonged Ca^2+^ elevations. *P*2*Y*-class receptors (mainly P2Y2) yield oscillatory Ca^2+^ transients.

Our simulations indicate that the activity of proteins belonging to the *G_αq_*/*IP*_3_ pathway determine whether *P*2*Y* receptors generate single, long duration peaks or oscillations. *P*2*Y* receptors are G protein coupled receptor (GPCR)s, of which P2Y2 promote *IP*_3_-dependent ER Ca^2+^ release via activating *G_αq_* proteins(42). This pathway includes PKC-dependent negative-feedback inhibition of *G_αq_*, which gives rise to stable *IP*_3_ oscillations and periodic intracellular Ca^2+^release (68). Negative feedback inhibition is a property of classical biochemical oscillators, for which the enzyme reaction rates determine the periodicity and decay of products like *IP*_3_(82), Although these oscillations may be stochastic, deterministic representations (82; 70) amenable to ordinary differential equation modeling are commonly used, given their ability to approximate the amplitude and peak-to-peak intervals of experimentally-measured Ca^2+^ release events (7). By modulating parameters like *kg_p_*_2*y*_, which controls the activation of *G_αq_*in our model, we identified how *IP*_3_ generation can be scaled to yield oscillatory versus single-peak waveforms commonly observed in microglia. Since these parameters represent the activity of proteins composing the *IP*_3_ synthesis pathways, waveforms exhibited in microglia are expected to be inherently sensitive to factors such as protein expression and co-localization (41).

Our simulations strongly implicate the role of ectonucleotidase (ENT) activity in controlling the responsiveness of microglia to extracellular ATP and related nucleotides. We demonstrate that neglecting ectonucleotidase activity in our model yielded sustained Ca^2+^ oscillations that were inconsistent with measurements in microglia cells and namely the data collected by Ikeda *et al* for MG5 microglial cells. This therefore suggests that ectonucleotidase activity determines the duration of ATP-mediated Ca^2+^ waveforms in microglia and ultimately cellular responses like migration and cytokine production. The ectonucleotidases CD39 and CD73 are the most highly expressed nucleotidases in microglia (50). CD39 rapidly hydrolyzes ATP and ADP into AMP, which curtailed Ca^2+^ waveforms within minutes of ATP treatment in our simulations. While our model of CD39 activity was parameterized to fit data from Robson *et al* (62), more detailed models such as from Sandefur *et al* (63) could give additional insights into how other expressed ectonucleotidases influence microglial responses to ATP. As an example, augmenting the CD39 model with contributions from the CD73 ectonucleotidase isoform(92), which metabolizes AMP into adenosine, will help determine which adenine metabolites predominate at the cell surface (61), as well as the receptors they stimulate.

Interestingly, it is increasingly recognized that extensions of the microglia plasma membrane infiltrate neural synapses which may be implicated in how glia scale and pruning neuron junctions Check this out: https://www.nature.com/articles/s41467-018-03566-5. ATP can be intermittently released in these junctions(62; 29), therefore we expect that the timescale and amplitude of those release events, as well as the rate by which ATP is metabolized, will control how microglia respond to these intercellular signals to fulfill their homeostatic functions. Here, spatially-explicit models of nucleotidases(61) that predict local ATP pools between interfaced cells could be important for determining how microglial responses in multi-cellular assemblies such as neural synapses differ from *in vitro preparations*.

### 5.3 Purinoreceptors control microglia migration and motility

Our study contributes a quantitative model linking P2Y12 activation to the PI3K and Akt axis that is essential for microglia migration (58; 36; 59). Unlike the metabotropic *P*2*Y* receptors implicated in intracellular Ca^2+^ signals, *P*2*Y* 12 activates *G_i/o_*, which directly stimulates PI3K and its phosphorylation of Akt. Our model reproduces the rate of PI3K-dependent Akt phosphorylation in addition to migration distances inferred from Ohsawa *et al* (58). Interestingly, both our model and data from Ohsawa *et al* (58) suggest that Akt phosphorylation is slow and reaches its maximum about three minutes after ATP treatment. This contrasts with the rapid onset of migration observed by others (19; 22). For instance, supplemental movies from Dou *et al* (19) indicate that microglia migrate almost immediately in response to ATP and approach a rate of [1.7 um/s] within 10 min of 1 mM ATP. Similar findings for microglia in tissue slices were also reported (22). Hence, either low levels of phosphorlylated Akt are sufficient for invoking migration at early timepoints, or alternatively, P2Y12- or *p*Akt-independent mechanisms mediate the rapid onset of migration. In support of the latter speculation, our model and experiments from Ohsawa *et al* (58) demonstrate that P2Y12 knock-out dramatically reduces, but does not eliminate, migration.

Given observations suggesting that 1) P2Y12 KO microglia migrate and 2) extracellular Ca^2+^ significantly enhances migration by promoting Akt phosphorylation (58; 59), our model was constructed to reflect the Ca^2+^-dependence of microglial migration. Importantly, our simulation results indicate that migration is significantly reduced when P2X contributions are neglected, in accordance with Ohsawa *et al* (58; 59). This finding suggests that there are ATP-triggered, Ca^2+^-dependent migration mechanisms (58; 36; 59) that could be sensitive to rapid Ca^2+^ signals, such as those exhibited by P2X4. These mechanisms could include Ca^2+^-dependent recruitment of PI3K to the plasma membrane (58), activation of the Ca^2+^-binding protein Iba (37), regulation of cytoskeletal proteins (48), and regulation of myosin by the CaM-dependent myosin light-chain kinase (66). To our knowledge, the rates of these mechanisms have not been examined in microglia, which precluded us from explicitly representing these processes in our model. However, we postulate that *P*2*X* receptors trigger Ca^2+^-dependent migration machinery that initiate migration, after which the gradual activation of the PI3K/Akt axis via *P*2*Y* 12 sustains migration over longer time intervals. Along these lines, low-amplitude Ca^2+^ oscillations from metabotropic P2Y receptors likely enhance migration, which is consistent with the requirement of *IP*_3_ induced Ca^2+^ release for P2Y12-driven chemotaxis (Kettenmann review (42). Clearly, ATP-induced migration in microglia is exceedingly complex (reviewed in (10; 73)) and warrants further investigation to unravel the intricate relationships between Ca^2+^ dynamics and migration.

Our simulations implicate Ca^2+^ signaling in promoting migration as well as TNF*α* synthesis. This raises the question as to whether ATP can stimulate microglia migration and motility associated with homeostatic functions ithout driving inflammatory cytokine responses. It is apparent from our simulations that a key distinction between these cellular responses is the duration of the intracellular Ca^2+^ waveform. Namely, our simulations show that submicromolar ATP treatments yield short-lived Ca^2+^ waveforms (<2 minutes) that are nonetheless sufficient for migration. In contrast, we show that higher amplitude Ca^2+^ waveforms or diminished CD39 activity are necessary for generating appreciable TNF*α* responses. Blocking ectonucleotidase activity prolongs *P*2*Y* and P2X4 Ca^2+^ transients and thereby increase TNF*α* mRNA production (Fig. S7). Similar prolonged Ca^2+^ signals are associated with inflammatory microglia (42; 34) and are routinely induced via millimolar ATP treatment *P*2*X*7, or reagents including LPS and ionomycin (34). It is apparent that the slow rate of activated transcription factor translocation into the nucleus which can occur over minutes (4) necessitates prolonged Ca^2+^ transients to induce transcription. This was reflected in our model for NFAT and was experimentally demonstrated for Ca^2+^ ionophore treated HEK293 cells in (4).

### 5.4 Differential purinergic receptor expression and its impact on microglia function

Our model suggests that the relative expression levels of P2 receptors enable microglia to regulate migration and pro-inflammatory responses to ATP (49). This occurs in part through modulating intracellular calcium dynamics. Since *P*2*X* and *P*2*Y* receptors exhibit unique and diverse Ca^2+^ waveforms (20), we hypothesized that phenotype-specific differences in P2 receptor expression in microglia influence both 1) Ca^2+^ responses and 2) migration. We investigated this hypothesis by adapting P2 expression levels in the model based on published BV2 cell mRNA data sets. Model predictions of ATP-stimulated Ca^2+^ waveforms and migration were compared against experiments with BV2 cells.

The mRNA data used for our model (30) indicated similar numbers of P2X4, P2X7, P2Y2 transcripts compared to primary cells. For simplicity, we assumed that the purinergic receptor activity in our model correlated with mRNA expression. However, data quantifying receptor expression and membrane localization is ultimately needed to accurately receptor activity. Nonetheless, based on our assumptions the model predicted Ca^2+^ waveforms in BV2 cells that were qualitatively similar to those simulated for primary cells. In contrast, P2Y12 mRNA transcripts were reduced 30-fold in BV2 cells relative to primary cells (30). Accordingly, our model predicted diminished migration in BV2 cells. We did not observe directed migration in our BV2 cell assays, which is consisted with other reports that confirm negligible or very slow migration responses to chemotactic stimuli (30; 24; 28) Nonetheless, we observed that the BV2 cells exhibited motility responses like membrane ruffling and membrane displacement in response to ATP. The motility processes increased in a ATP-dose dependent manner consistent with our model predictions for migration.

## 6 Limitations

There are several model limitations that can guide refinement in subsequent studies. A prominent limitation is that many of the underlying biological processes linking ATP binding to migration and cytokine responses are not completely resolved. Of those we considered in our model, the kinetics of those processes are also imprecise. The Ca^2+^ responses induced by the purinoreceptors are perhaps the best characterized of these processes, as time-dependent fluorescence data were available. Other processes though were heavily reliant on western blotting and microscopy, which are much less precise. We also assumed that the biochemical pathways are spatially homogeneous within the cell for the simplicity of modeling and parameterization. Nonetheless, a number of proteins have precise subcellular localization or undergo changes in a localization about activation, such as P2X4(78) and NFAT(4). Accounting for these changes could impact their ability to promote gene transcription versus motility or migration responses. Along these lines, for simplicity we considered only changes in P2 mRNA expression when extending our model to the BV2 microglial cells. Significant changes in other downstream proteins mediating migration responses would be expected to have an impact on migration.

It was evident from our measurements of ATP-induced Ca^2+^ waveforms that cell-to-cell variation in responses was substantial. Namely, many of the individual cells presented traces that strongly deviated from the mean (see Fig. 7) Our modeling approach relies on deterministic equations, which are most appropriate for describing the average behavior of a large ensemble of cells. This approach is valid, given that many of the experimental data used to train our approach were from western blots and mRNA quantification, which generally use large pools of cells. Stochastic models such as that from Skupin or Cao *et al* (6; 68; 67) could be used in complement to our model to investigate how cell-to-cell variations in gene expression or protein activity could impact ATP-induced Ca^2+^ waveforms.

Similarly, our experimental measurements of BV2 yielded largely non-oscillatory waveforms. Nonetheless, stimulation of BV2 cells with ATP yielded oscillatory Ca^2+^ transients in a small subset of cells. This raises the possibility that this subset expresses *P*2*Y* receptors at a greater level than the population average. Cell-to-cell variations in purinergic receptor transcripts or expression have not been characterized in BV2 cell lines, which limits our ability to associate oscillations in subsets of BV2 cells with P2X or P2Y activity. However, several single-cell RNAseq studies(14; 15; 30) have been conducted for primary microglia cultures, which indicate there exist sub-populations with unique patterns of *P*2*X* versus *P*2*Y* receptor expression. At the very least, the cell-to-cell variability underscores a need for single-cell characterization of cell genotypes and phenotypes, as well as sensitivity analyses such as in Fig. 6 to better characterize Ca^2+^ waveforms and their effects in diverse microglial cell populations.

Lastly, our model could be improved by accounting for *K*^+^-dependent signaling in particular. There are a multidude of mechanisms by which changes in intercellular *K*^+^ and membrane potential could influence microglial signal transduction, such as by enhancing the electromotive force for Ca^2+^ entry, or by influencing the activity of the sodium/*K*^+^ ATPase. Reflecting these contributions may provide more complete descriptions of *K*^+^ mechanisms mediating inflammation (72) and migration (76). For instance, *P*2*Y* 12-dependent activation of *K*^+^ channel dynamics is believed to contribute to migration (42). This is supported by studies suggesting that *P*2*Y* 12-activation induces substantial outward current associated with *K*^+^ channel activity (76; 23). It is further understood that *K*^+^ efflux constitutes an important stage of priming the microglial inflammasome, which is necessary for maturating pro-inflammatory cytokines such as IL-1*β* (84). Along these lines, it is increasingly recognized that P2X7 and P2X4 conduct *K*^+^ countercurrent when activated (55; 87); moreover, changes in *K*^+^-channel expression upon microglial activation may contribute to these responses (54).

While we modeled PI3K, RAGE/RhoaA/ROCK are also involved in mediating chemotaxis(77; 90). For instance, it was shown that inhibition of ROCK via H-1152 reduced p38 phosphorylation and membrane ruffling following ATP-dependent P2Y12/13 activation (77). p38 is known also to be Ca^2+^sensitive (80), which again introduces aother potential pathway sensitive to Ca^2+^ waveform characteristics. Inhibition of MLCK, Rac1 and p38-MAPK also results in the attenuation of motility in primary cultured murine microglia(51). CaM dependent contributions to migration are also important to recognize. O’Brien *et al* found that active CaM was involved in chemotaxis by monitoring the activity of its target, phosphodiesterase (PDE1) (57). Similarly, Yao *et al* demonstrated the involvement of CaM in migration via CaM-dependent myosin light chain kinase (MLCK)(86). Altogether, the dependencies of cell migration on diverse signaling pathways suggest microglia are highly adaptive to a variety of extracellular stimuli to promote motility responses. Hence, resolving these inter-dependencies will warrant additional studies and their dependency on Ca^2+^ to delineate microglial migration responses specific to ATP.

## 7 Conclusions

ATP-induced Ca^2+^ waveforms in microglia have diverse properties, such as amplitude, duration and oscillatory behavior. These properties depend on which P2 receptor types are activated, in addition to ectonucleotidase activity. In this study, we developed a computational model to predict how P2 receptors and ectonucleotidase hydrolases control Ca^2+^ waveforms in microglia that in turn influence microglia migration and cytokine production. With this model, we examine the propensity for these diverse Ca^2+^ waveforms to drive these canonical microglial responses to ATP.

We interpret these results in light of our previously published microglia model for probing P2X contributions to TNFa mRNA synthesis as a model for pro-inflammatory cytokine responses to ATP (9). In that study, we demonstrated that Ca^2+^ waveforms generated by P2X activation are typically high-amplitude and of finite duration. With the addition of *P*2*Y*, we predict a wider diversity of Ca^2+^ waveforms that can include stable and damped oscillations of low amplitude and frequency. Interestingly, our modeling results highlight a complementary role of ectonucleotidase activity, namely CD39, in controlling the ATP pool available to trigger such responses, by hydrolyzing ATP and ADP into AMP. This finding is of particular importance, given that CD39 and a complementary ectonucleotidase, CD73, of which the latter hydrolyzes AMP and AMP into adenine, are highly expressed in microglia and undergo significant changes in expression (39).

Our simulations indicate that the distinct Ca^2+^ waveforms shaped by P2 receptors and CD39 selectivity control downfield signaling pathways. We investigated this selective control by simulating migration versus cytokine production responses as a function the Ca^2+^ waveforms generated by P2 receptors. Our model indicates that short-duration, oscillatory Ca^2+^ transients induced by P2Y receptors and P2X4 with micromolar ATP are sufficient to promote migration responses without significantly inducing TNFa production. On the other hand, millimolar ATP concentrations that activated P2X7 supported sustained cytosolic Ca^2+^ levels that could trigger TNF*α* release. We speculate that these findings illustrate how microglia orchestrate complex cytokine and migration functions in response to ATP as well as other damage associated molecular patterns.

The concerted roles of P2X, P2Y and ectonucleotidase proteins in mediating cellular responses to ATP further suggest how changes in gene expression shape microglia responses to stimuli. For instance, higher P2Y12 expression in resting relative to pro-inflammatory microglia likely favor migration responses to ATP in the former. Similarly, elevated P2X4 and P2X7 expression in proinflammatory microglia sensitize cytokine responses to ATP. These differences in microglia responses following changes in gene programming underscore the need for models to account for changes in protein activity.

Robust characterization of detailed signaling networks in diverse microglia phenotypes remains a significant challenge. This is especially challenging for tissue resident microglia that are difficult to experimentally manipulate *in situ*. For this reason, we tested if our model could leverage mRNA transcript data from the BV2 microglial cell line to approximate changes in P2 receptor activity. Using those mRNA data, we found that the model predicted Ca^2+^ waveforms and migration responses to ATP that were reasonably consistent with experiments we conducted using the BV2 microglia cell line. This raises the possibility that coupling models trained from cultured primary or immortalized cells in vitro with transcriptomic and proteomic data could enable predictions of microglia behavior *in vivo*, where extensive functional testing is not feasible. Related to this, since dysfunctional microglial responses are associated with neurological disorders including chronic pain, Alzheimer’s Disease and Parkinson’s Diseases (12), our computational model may be an invaluable tool to probe mechanisms underlying these diseases.

## 8 Acknowledgements

Research reported in this publication was supported by the Maximizing Investigators’ Research Award (MIRA) (R35) from the National Institute of General Medical Sciences (NIGMS) of the National Institutes of Health (NIH) under grant number R35GM124977. This work used the Extreme Science and Engineering Discovery Environment (XSEDE)(79), which is supported by National Science Foundation grant number ACI-1548562. All the figures are processed or adapted from BioRender.com.

## 9 Supplement

### S.1 Tables

**Table S1:** Reactions used in the computational microglia model.

| Rxn No. | Rxn | Cell | Reference |
| --- | --- | --- | --- |
| 1 | $P2Y \Rightarrow IP_3 \Rightarrow [Ca^{2+}]_i$ | heptaocyte | (16) |
| 2 | $P2X4 \Rightarrow [Ca^{2+}]_i$ | microglia | (78; 26) |
| 3 | $P2X7 \Rightarrow [Ca^{2+}]_i$ | microglia | (44; 8) |
| 4 | $[Ca^{2+}]_e \Rightarrow [Ca^{2+}]_i$ (leak) | | Fit |
| 5 | $NCX \Rightarrow [Ca^{2+}]_e$ | microglia | (65) |
| 6 | $[Ca^{2+}]_i \Rightarrow$ Buffers | | Fit |
| 7 | $[Ca^{2+}]_i \Rightarrow CaM \Rightarrow CN$ | Cardiac | (31; 4) |
| 8 | $[Ca^{2+}]_i \Rightarrow SERCA$ | Cardiac | (65) |
| 9 | $[Ca^{2+}]_{ER} \Rightarrow [Ca^{2+}]_i$ (leak) | | Fit |
| 10 | $[Ca^{2+}]_{ER} \Rightarrow$ Calreticulin (Calsequestrin) | astrocytes | (65) |
| 11a | $[Ca^{2+}]_i \Rightarrow$ p-p38 | microglia | (80) |
| 11b | $p-p38 \Rightarrow TNF\alpha$ | microglia | (33) |
| 12a | $CN \Rightarrow NFAT$ | microglia | (88; 13) |
| 12b | NFAT cycle | myocyte | (13) |
| 12c | $NFAT \Rightarrow TNF\alpha$ | microglia | (53) |
| 13 | $TNF\alpha$ Production | microglia | Fit |
| 14 | Activation of $P2X7 \Rightarrow TNF\alpha$ release | microglia | (3) |
| 15 | $P2Y12 \Rightarrow G_{i/o}$ | microglia | (42; 59) |
| 16,17 | $G_{i/o} \Rightarrow PI3K \Rightarrow pAkt$ | microglia | (59; 23) |
| 18 | $pAkt + [Ca^{2+}]_i \Rightarrow$ Migration | microglia | (59) |
| 19 | NTPDase of ATP | COS-7 | (47; 62) |

**Table S2:**
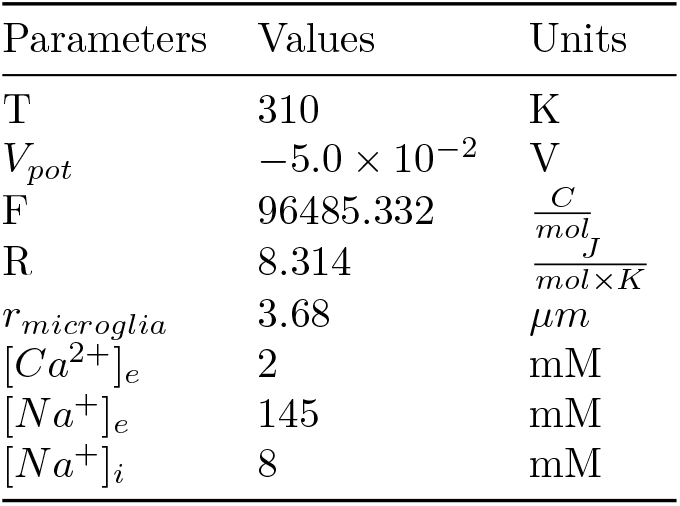
General Parameters

**Table S3:** Parameters associated with S.2.2

| Parameters | Values | Units |
| --- | --- | --- |
| $kd_{P2Y}$ | 410.0 | nM/s |
| $hd_{P2Y}$ | 4.0 | 1/s |
| $ld_{P2Y}$ | 0.3 | nM/s |
| $kc_{P2Y}$ | $1.5 \times 10^6$ | nM/s |
| $k_{rec,P2Y}$ | 0.01 | 1/s |
| $h_{rec,P2Y}$ | 10.0 | 1/s |
| $Kp_{P2Y}$ | 20.0 | 1/s |
| $Kc_{P2Y}$ | 100.0 | 1/s |
| $Kg_{P2Y}$ | 50.0 | nM |
| $Kd_{P2Y}$ | 5.0 | nM |
| $Ks_{P2Y}$ | 5.0 | nM |
| $[PH]$ | 0.0 | nM |
| $ka_{P2Y}$ | 1000.0 | nM |
| $n_{Ca^{2+},P2Y}$ | 3.0 | |
| $[P2Y]_{total}$ | 200.0 | nM |
| $G_{\alpha,total}$ | 200.0 | nM |
| $n_{P2Y}$ | 2.0 | |
| $m_{P2Y}$ | 4.0 | |
| $Kd_{ATP,P2Y}$ | 250.0 | μM |
| $K_{deg,P2Y}$ | 0.1 | 1/s |

**Table S4:** Parameters associated with *P*2*X*4 receptor kinetics (S.2.3)

| Parameters | Values | Units |
| --- | --- | --- |
| $k_1$ | 1.0 | 1/s |
| $k_2$ | $2.61 \times 10^5$ | 1/(M×s) |
| $k_3$ | 0.1 | 1/s |
| $k_4$ | $1.6 \times 10^5$ | 1/(M×s) |
| $k_5$ | 0.25 | 1/s |
| $k_6$ | $8.0 \times 10^6$ | 1/(M×s) |
| $H_1$ | 0.02 | 1/s |
| $H_2$ | 0.0 | 1/s |
| $H_6$ | 0.1 | 1/s |

**Table S5:** Parameters associated with *P*2*X*7 receptor kinetics (S.2.3)

| Parameters | Values | Units |
| --- | --- | --- |
| $k_1$ | 190 | 1/s |
| $k_2$ | $8.13 \times 10^3$ | 1/(M×s) |
| $k_3$ | 0.04 | 1/s |
| $k_5$ | 0.07 | 1/s |
| $H_1$ | $5.0 \times 10^{-3}$ | 1/s |
| $H_2$ | 0.3 | 1/s |
| $H_5$ | 0.0 | 1/s |
| $H_6$ | 0.0 | 1/s |
| $k_{4,low}$ | $1.0 \times 10^2$ | 1/(M×s) |
| $k_{6,low}$ | $5.0 \times 10^2$ | 1/(M×s) |
| $H_{7,low}$ | $1.0 \times 10^3$ | 1/s |
| $k_{4,high}$ | $7.0 \times 10^3$ | 1/(M×s) |
| $k_{6,high}$ | 0.1 | 1/(M×s) |
| $H_{7,high}$ | 0.008 | 1/s |
| $k_d$ | 420 | $\mu M$ |
| $n$ | 15 | |

**Table S6:** Parameters associated with the estimation of inward current and corresponding Ca^2+^ influx (S.2.3)

| Parameters | Values | Units |
| --- | --- | --- |
| $G_{12,P2X4}$ | $6.15 \times 10^{-10}$ | $\frac{C}{s \times V}$ |
| $G_{12,P2X7}$ | $1.0 \times 10^{-8}$ | $\frac{C}{s \times V}$ |
| $E_{12,P2X4}$ | 0.0 | V |
| $E_{12,P2X7}$ | 0.0 | V |
| $f_{I_{Ca^{2+}},P2X4}$ | 0.0824 | |
| $f_{I_{Ca^{2+}},P2X7}$ | 0.1 | |
| $f_{conv.,P2X4}$ | 11 | |
| $f_{conv.,P2X7}$ | 1 | |

**Table S7:** Parameters for NCX mechanisms associated with S.2.4

| Parameters | Values | Units |
| --- | --- | --- |
| $Q_{10}$ | 1.20 | |
| $Kd_{Act}$ | 40.0 | nM |
| $n_H$ | 3.44 | |
| $H_{Na}$ | 3.60 | |
| $V_{max}$ | 35 | A/F |
| $\eta$ | 0.70 | |
| $k_{sat}$ | 0.04 | |
| $K_{max,[Ca^{2+}]_i}$ | $3.63 \times 10^3$ | nM |
| $K_{max,[Na^+]_i}$ | $1.23 \times 10^7$ | nM |
| $K_{max,[Na^+]_e}$ | $8.75 \times 10^7$ | nM |
| $K_{max,[Ca^{2+}]_e}$ | $1.30 \times 10^6$ | nM |
| $C_{mem}$ | $1.2 \times 10^{-11}$ | F |

**Table S8:** Parameters for SERCA mechanisms associated with S.2.4

| Parameters | Values | Units |
| --- | --- | --- |
| $Q_{10}$ | 2.6 | |
| $V_{max}$ | $9.09 \times 10^6$ | nM/s |
| $K_f$ | $2.80 \times 10^2$ | nM |
| $K_r$ | $2.10 \times 10^6$ | nM |
| $H$ | 1.787 | |

**Table S9:** Parameters for CaM/CN and NFAT cycle calculations shown in S.2.7

| Parameters | Values | Units |
| --- | --- | --- |
| $k_{ab}$ | $1.0 \times 10^{-5}$ | $1/(nM^2 \times s)$ |
| $k_{ba}$ | 10.0 | 1/s |
| $k_{bc}$ | $1.0 \times 10^{-4}$ | $1/(nM^2 \times s)$ |
| $k_{cb}$ | $1.0 \times 10^3$ | 1/s |
| $k_{on,A}$ | $1.0 \times 10^{-2}$ | $1/(nM \times s)$ |
| $k_{off,A}$ | 1.0 | 1/s |
| $k_{on,B}$ | $2.0 \times 10^{-6}$ | $1/(nM^2 \times s)$ |
| $k_{off,B}$ | 1.0 | 1/s |
| $[CaM]_{total}$ | 100 | nM |
| $[CN]_{total}$ | 67 | nM |

**Table S10:** Parameters for phosphorylation of p38 and Ca^2+^ buffer calculations listed in S.2.6

| Parameters | Values | Units |
| --- | --- | --- |
| $[pp38]_{total}$ | 100 | |
| $k_{b,pp38}$ | $8.51 \times 10^{-4}$ | 1/s |
| $k_{f,pp38}$ | $1.1 \times 10^{-2}$ | 1/s |
| $k_{d,pp38}$ | 150 | nM |
| $n_{pp38}$ | 5 | |
| $B_{max,F}$ | $2.5 \times 10^4$ | nM |
| $k_{on,F}$ | 0.15 | 1/(nM×s) |
| $k_{off,F}$ | 23.0 | 1/s |
| $B_{max,B}$ | $1.0 \times 10^4$ | nM |
| $k_{on,B}$ | 1.0 | 1/(nM×s) |
| $k_{off,B}$ | $1.0 \times 10^3$ | 1/s |
| $B_{max,S}$ | $1.4 \times 10^5$ | nM |
| $k_{on,S}$ | 0.1 | 1/(nM×s) |
| $k_{off,S}$ | $6.5 \times 10^4$ | 1/s |

**Table S11:** Parameters for simulating TNF*α* synthesis and its exocytosis listed in S.2.8

| Parameters | Values | Units |
| --- | --- | --- |
| $k_{transcript}$ | $2.78 \times 10^{-4}$ | 1/s |
| $k_{trnsl}$ | $2.0 \times 10^{-4}$ | 1/s |
| $k_{deg,TNF\alpha}$ | $1.38 \times 10^{-2}$ | 1/s |
| $k_{deg,mRNA}$ | $1.35 \times 10^{-4}$ | 1/s |
| $IC50_1$ | 0.4 | |
| $n_1$ | 2 | |
| $IC50_2$ | 75.0 | |
| $n_2$ | 5.5 | |
| $k_{exp,f}$ | $5.11 \times 10^{-4}$ | 1/(molecule×s) |
| $k_{exp,r}$ | $1.78 \times 10^{-4}$ | 1/s |
| $D_{nc}$ | 10.0 | 1/s |
| $D_{exo}$ | 5.0 | 1/s |
| $k_d$ | 25.0 | nM |

**Table S12:** Parameters for simulating TNF*α* synthesis and its exocytosis listed in S.2.9

| Parameters | Values | Units |
| --- | --- | --- |
| $k_{f,1}$ | 0.008 | 1/s |
| $k_{b,1}$ | 0.02 | 1/s |
| $k_{f,2}$ | 0.1 | 1/s |
| $k_{b,2}$ | 0.01 | 1/s |
| $k_{f,3}$ | 0.01 | 1/s |
| $k_{b,3}$ | 0.01 | 1/s |
| $k_{f,4}$ | 0.00001 | 1/s |
| $k_{b,4}$ | 0.01 | 1/s |
| $k_{f,5}$ | 0.001 | 1/s |
| $k_{b,5}$ | 0.1 | 1/s |
| $k_{f,6}$ | 0.001 | 1/s |
| $k_{b,6}$ | 0.01 | 1/s |
| $k_{deg,1}$ | 0.05 | 1/s |
| $[P2Y12]_{total}$ | 100 | |
| $[PI3K]_{total}$ | 100 | |
| $[Akt]_{total}$ | 100 | |

**Table S13:** Parameters for the degradation of ATP by NTPDase listed in S.2.12

| Parameters | Values | Units |
| --- | --- | --- |
| $k_{1,deg}$ | 0.002 | 1/s |
| $k_{2,deg}$ | 0.008 | 1/s |

### S.2 Methods

#### S.2.1 Model equations

#### S.2.2 GPCR-model: Model 5 - Cuthbertson and Chay

These equations are implemented and integrated with the mathematical expression for the *P*2*X*-mediated Ca^2+^ dynamics

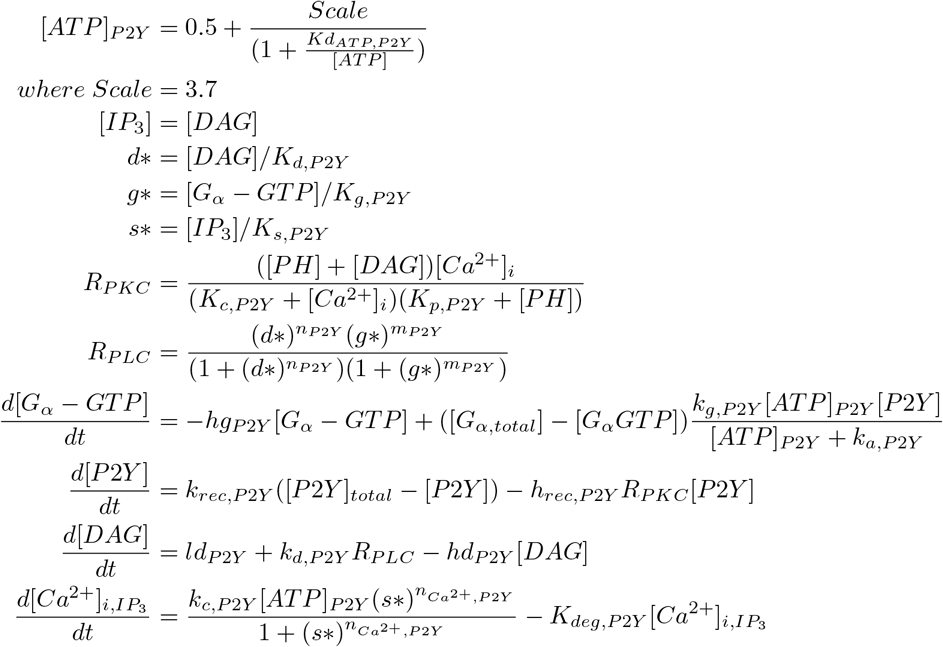

#### S.2.3 P2X4 kinetic models - the model was taken from others but implemented

##### *P*2*X*4 Dynamics

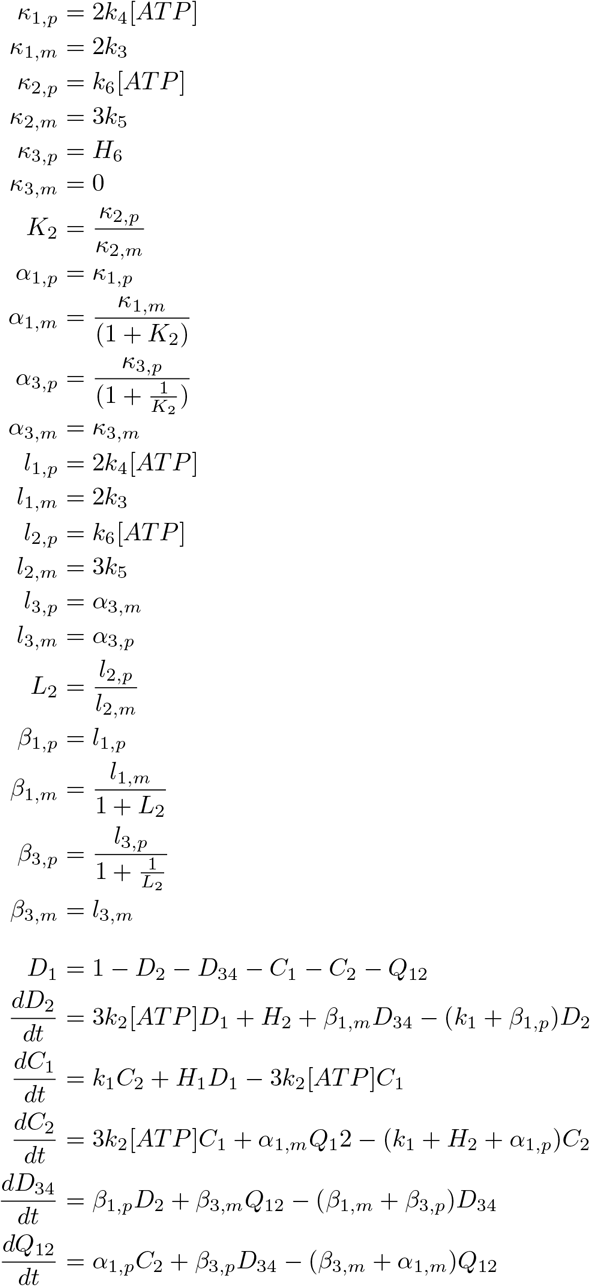

##### *P*2*X*7 Dynamics

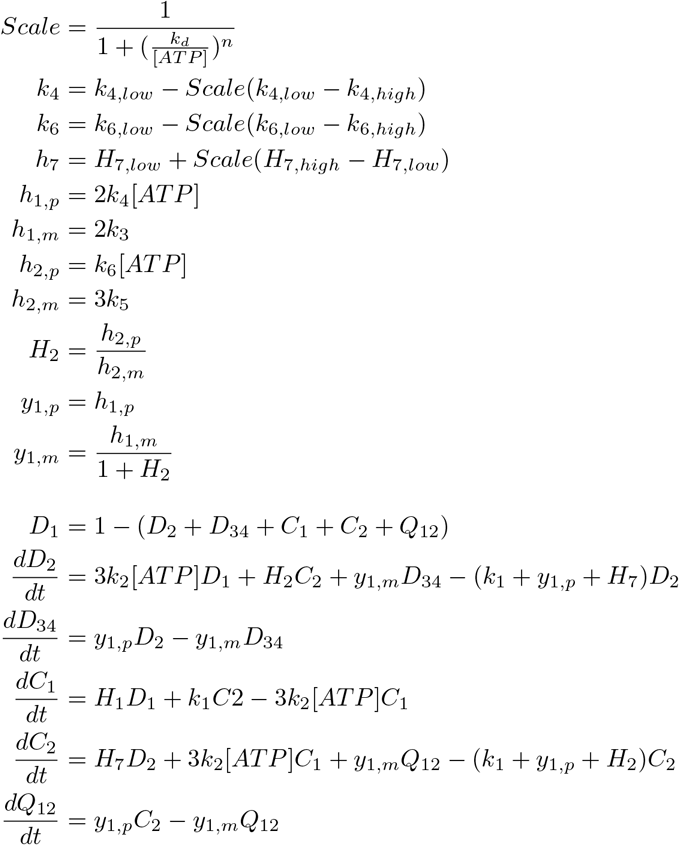

##### Estimation of P2X-induced inward current and Ca^2+^ influx

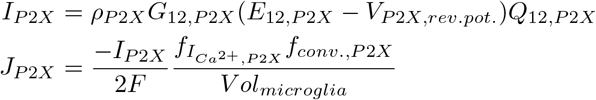

#### S.2.4 NCX and SERCA model - Shannon-Bers model

##### NCX

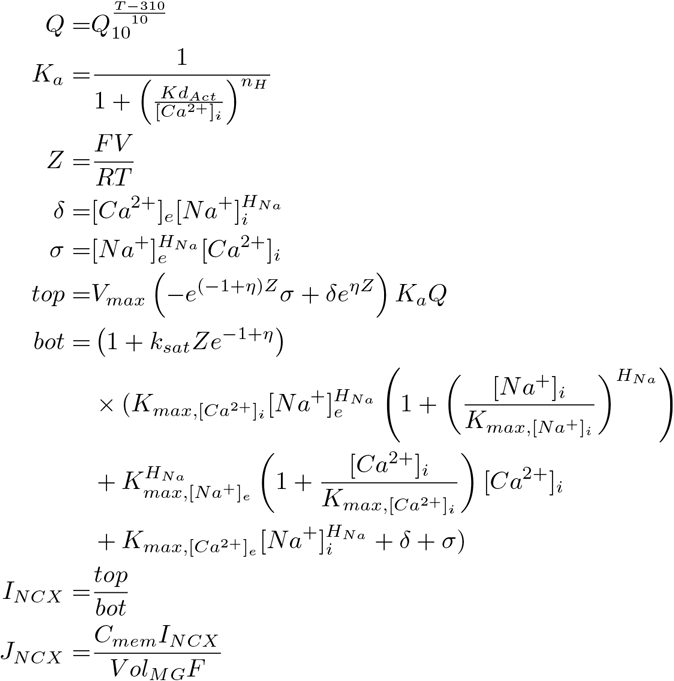

##### SERCA

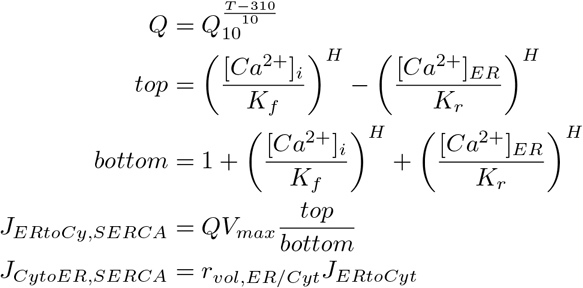

#### S.2.5 Leak terms

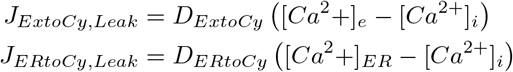

#### S.2.6 Phosphorylation of p38 and buffers

F,S, and B denote Fura-2, Calsequestrine in ER, and Extra unknown buffer in cytoplasm

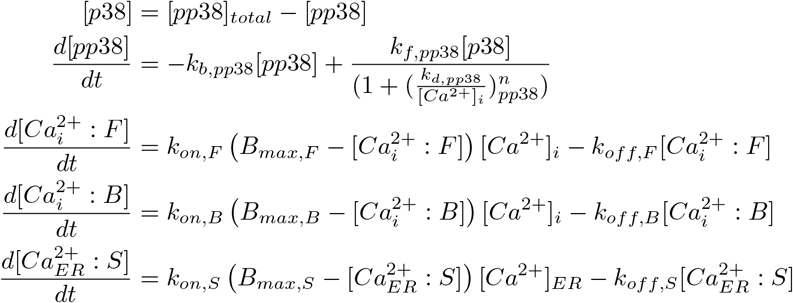

#### S.2.7 CaM/CN and NFAT cycle

##### Chemical Reaction Equations

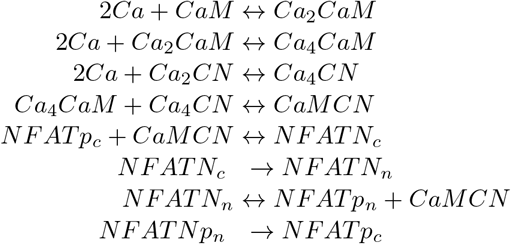

##### CaM/CM Activation

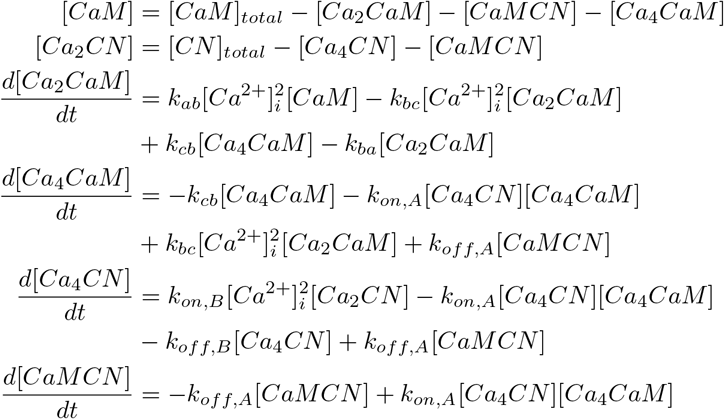

##### NFAT Cycle

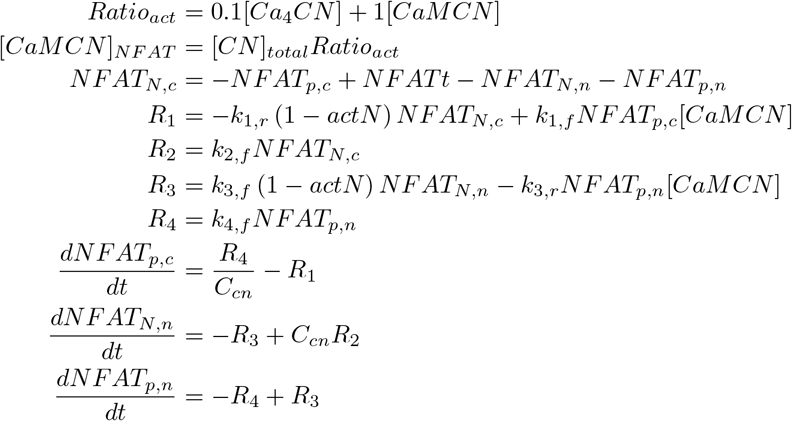

*C_cn_* is the volume fraction between cytosol and nucleus

#### S.2.8 TNF*α* synthesis

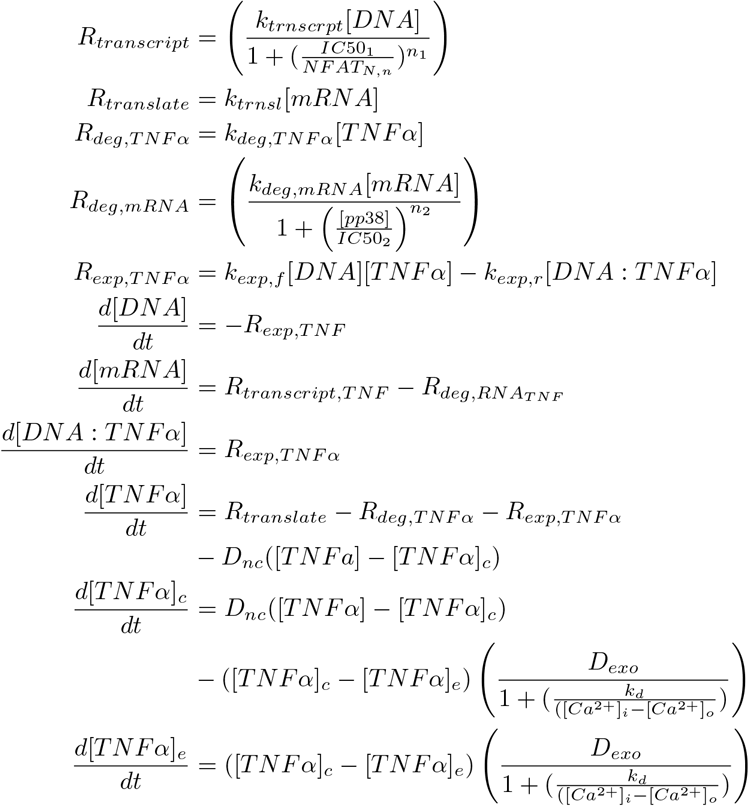

#### S.2.9 *P*2*Y* 12-mediated Signaling and Chemotaxis

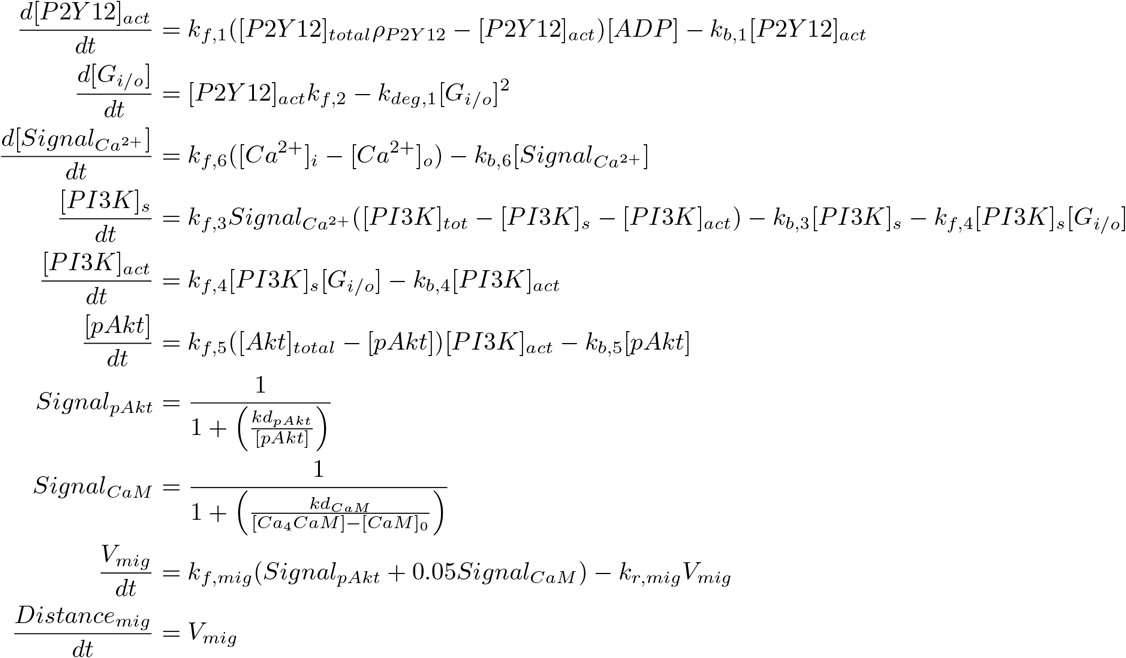

#### S.2.10 [*Ca*^2+^]_*ER*_ Homeostasis Equations

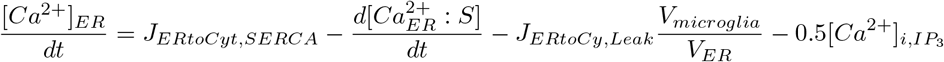

#### S.2.11 [*Ca*^2+^]_*i*_ Homeostasis Equations

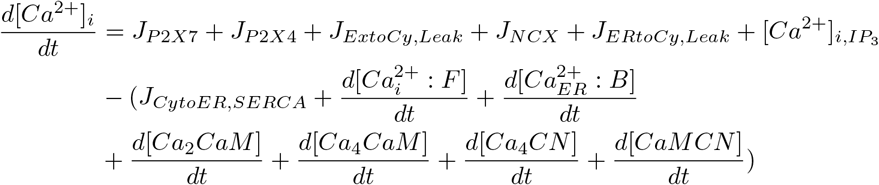

#### S.2.12 Degradation of ATP by NTPDase1

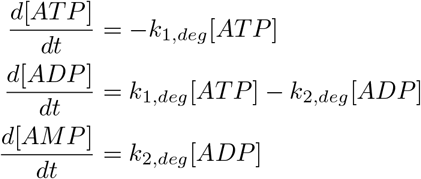

### S.3 How data was analyzed or Genetic Algorithm

Fitting was done according to our genetic algorithm in (9). State variable values as a function of time were plotted in jupyter notebooks for analysis. The current version of the genetic algorithm we implemented in this work generates multiple child parameters based on the initial guess. The algorithm also takes multiple observable to prevent the parameter set from being limited to a single reference data. Lastly, this version allows us to accelerate the fitting by processing multiple parameters simultaneously.

### S.4 Figures

**Figure S1:**
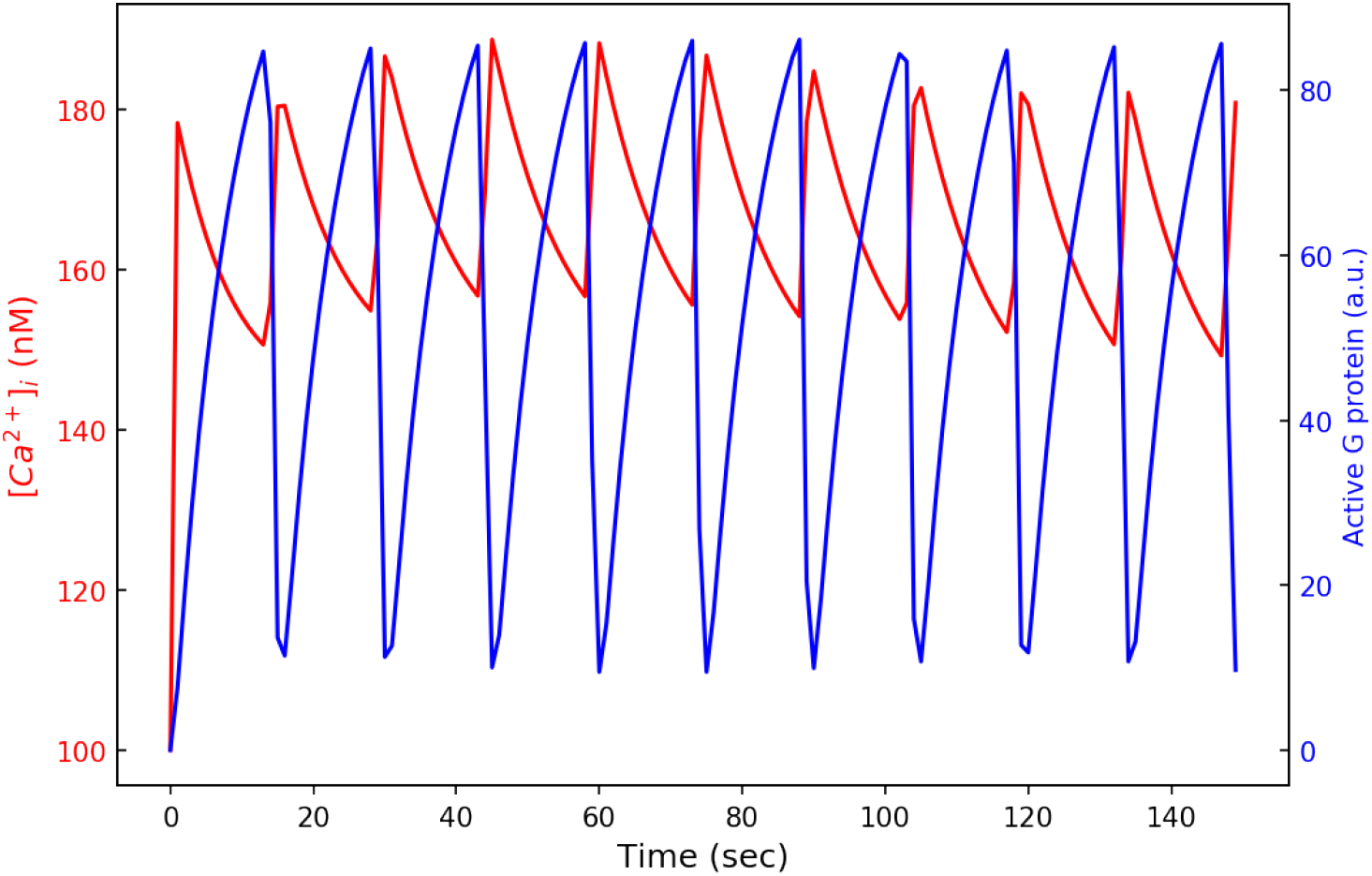
The demonstration of Ca^2+^ fluctuation mediated by P2Y-class receptor activation and its synchronization with the active *G_αq_*. .

**Figure S2:**
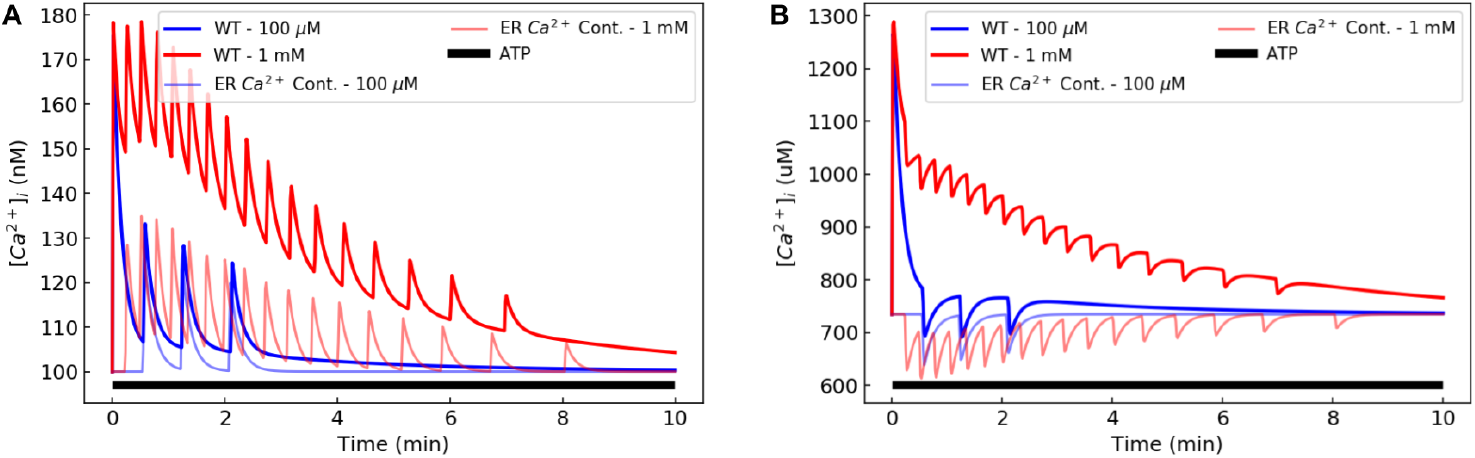
Comparison between Ca^2+^ transients induced by ER Ca^2+^ release via *IP*_3_-mediated pathway at 100 μM and 1 mM ATP concentrations in cytoplasm (A) and ER lumen (B). The WT microglia model was used for this prediction. The faded lines denote the contribution by *P*2*Y* receptor activation that results in ER Ca^2+^ release to the cytosolic domain. The data demonstrate the relationship between cytosolic and ER Ca^2+^ transients, which suggest that roughly 43.7% of Ca^2+^ is drawn from the ER at low ATP vs. 33.3% at high ATP. .

**Figure S3:**
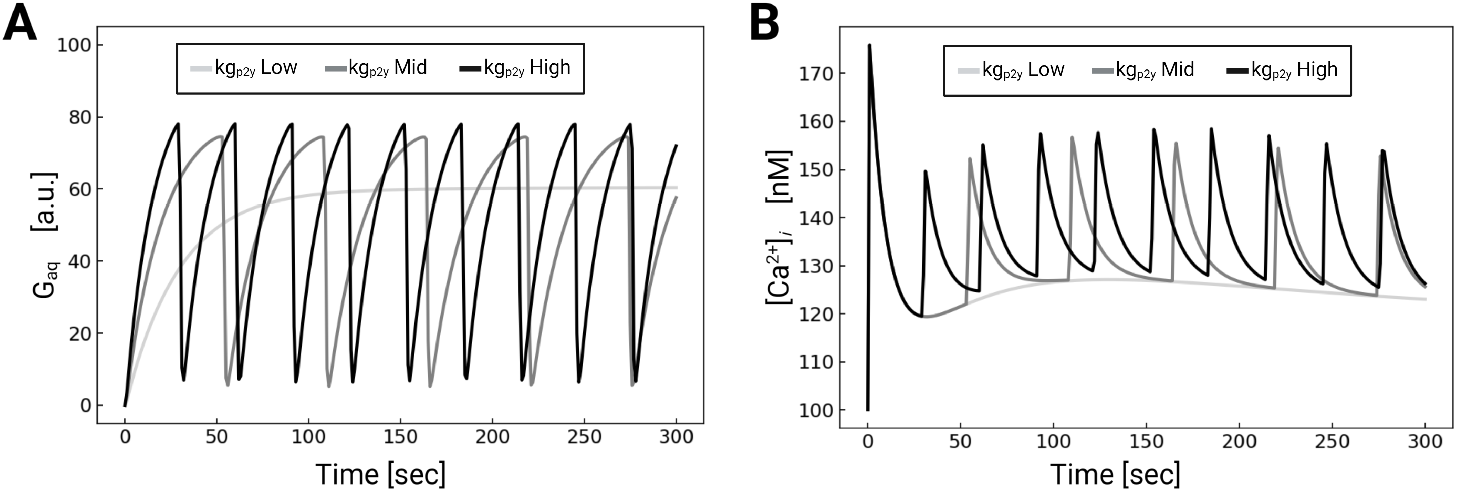
Oscillation of active *G_αq_* and their corresponding intracellular Ca^2+^ transients with respect to *kg_p_*_2*y*_ that controls the activation of *G_αq_*. .

**Figure S4:**
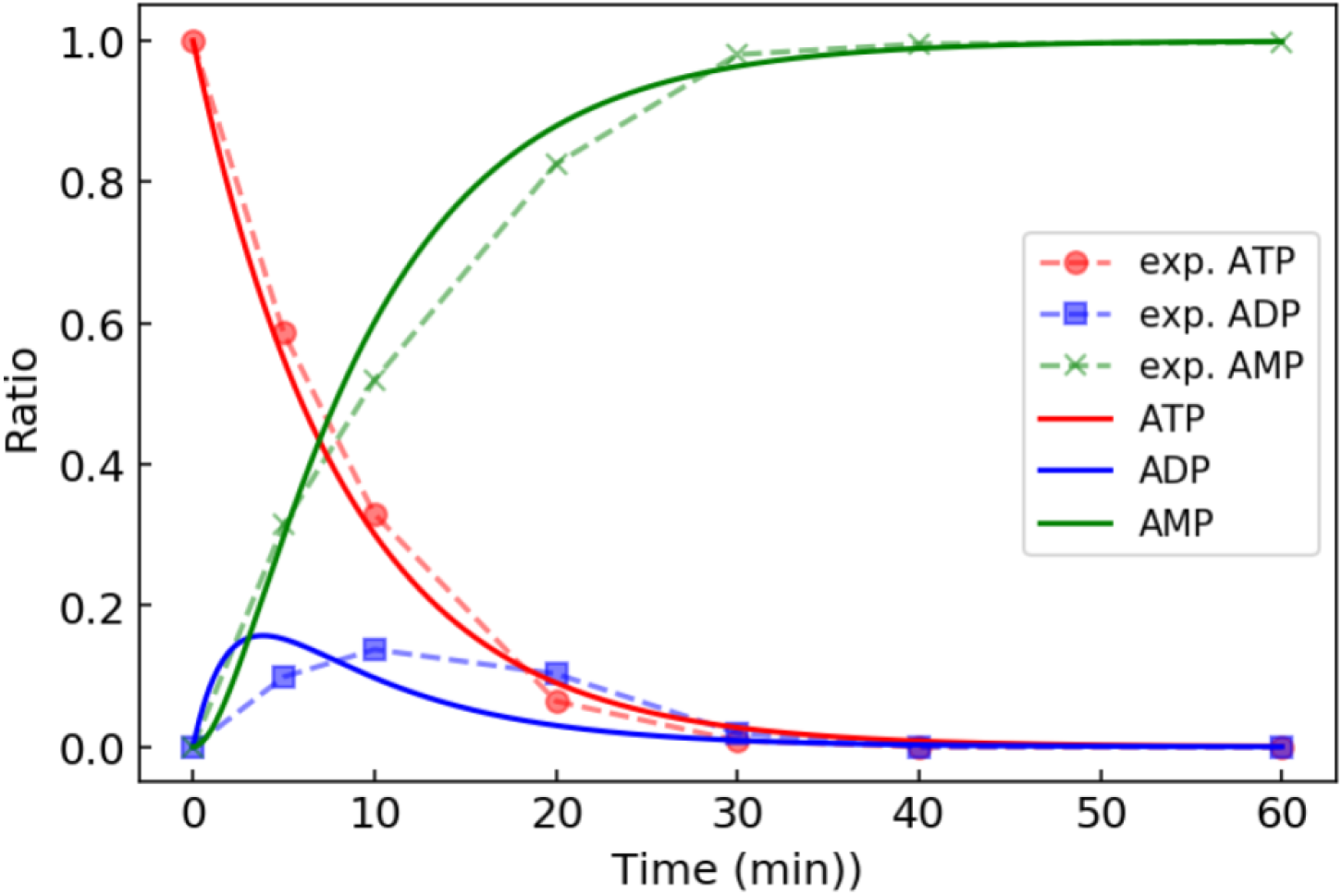
Validations of our model of ATP/ADP hydrolysis into AMP by CD39 against experimental data (dashed). Each nucleotide concentration was measured by Kukulski *et al* (47) in COS-7 cells.

**Figure S5:**
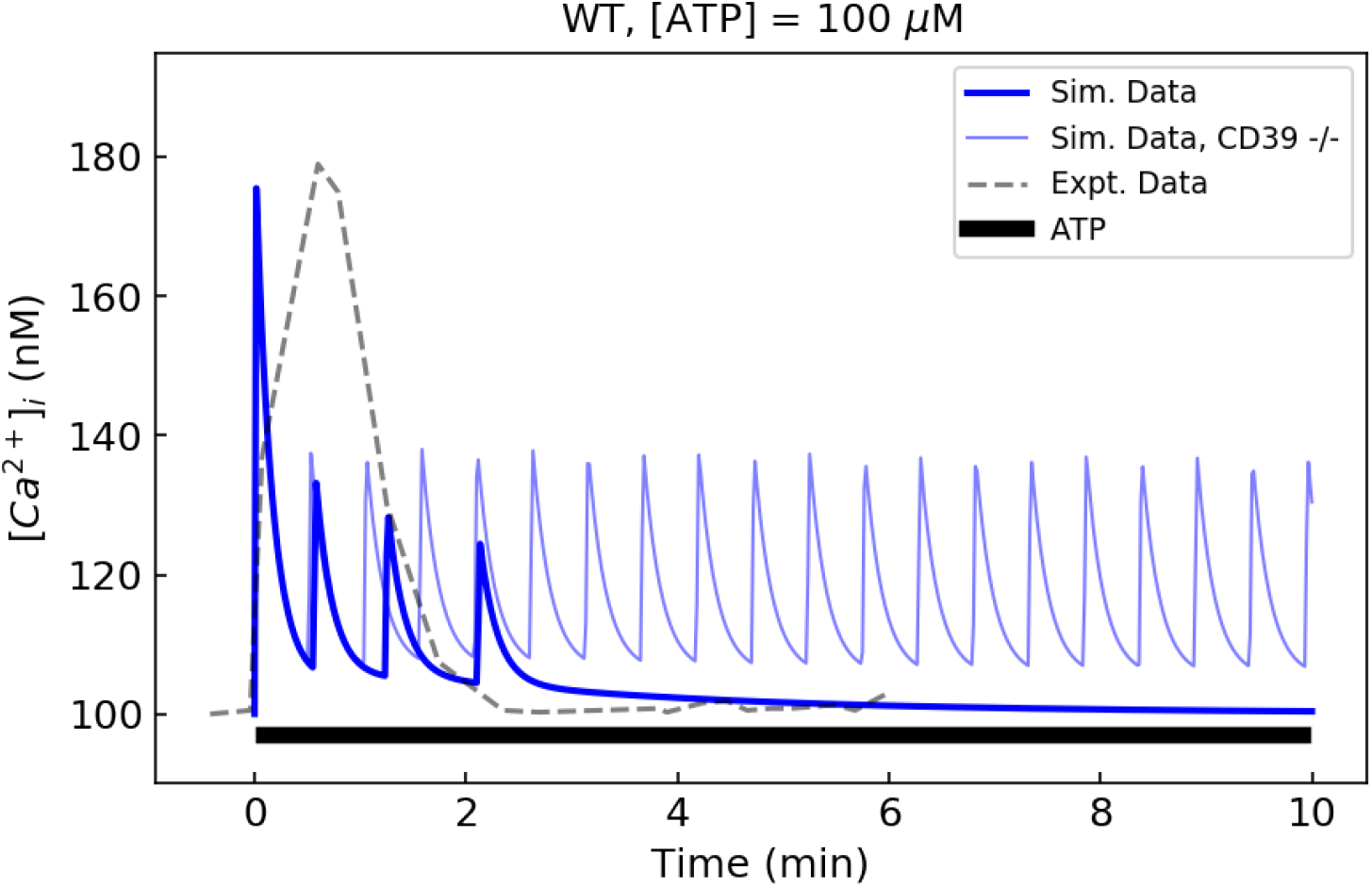
100 μM ATP treatment preferentially activates *P*2*X*4 receptors and yields a similar Ca^2+^ transient to that of *P*2*X*7. In both experiment and simulation, low ATP stimulation (100 μM) applied to WT microglia was sufficient to activate all purinergic receptors except for *P*2*X*7. .

**Figure S6:**
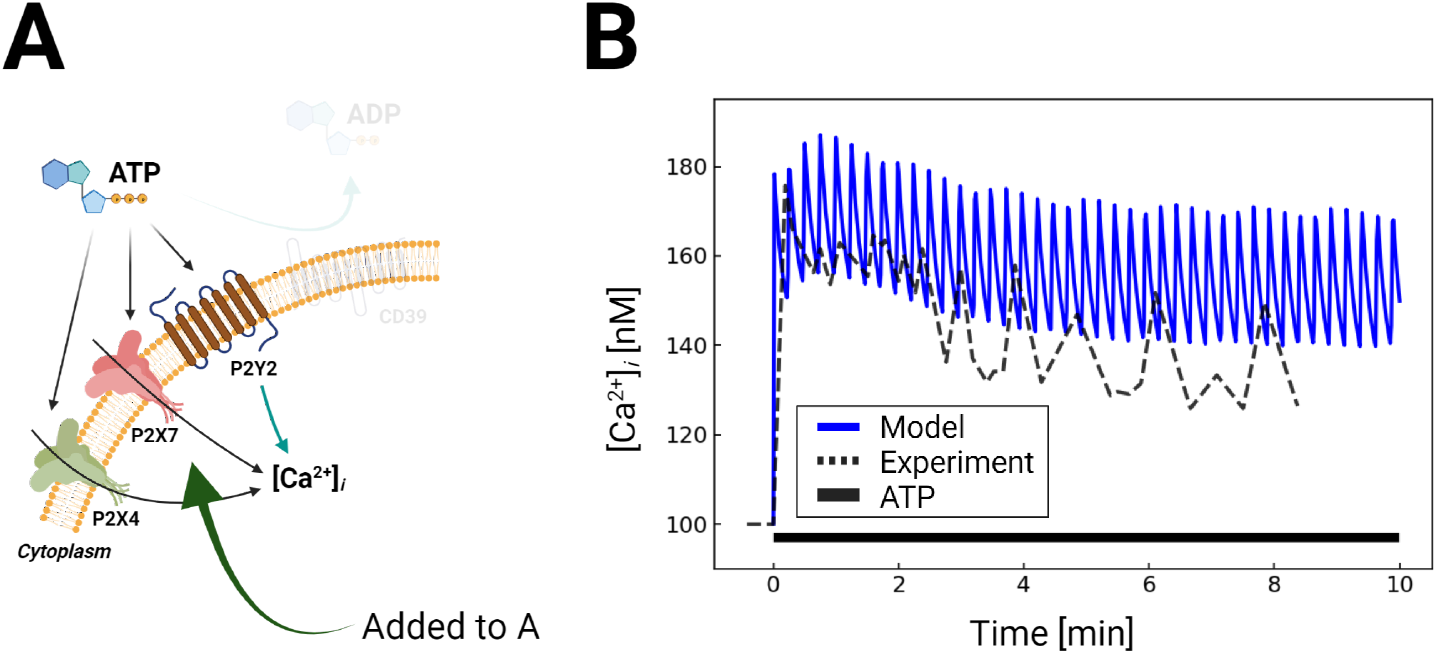
A) Schematic for Ca^2+^ waveforms generated in a control system comprising P2X4, P2X7 and P2Y2 in response to 1mM ATP for 5 minutes, but exclusing CD39 nucleotidase activity. B) Comparison of predicted (blue) and experimentally-measured (35) (dashed) Ca^2+^ waveforms. .

**Figure S7:**
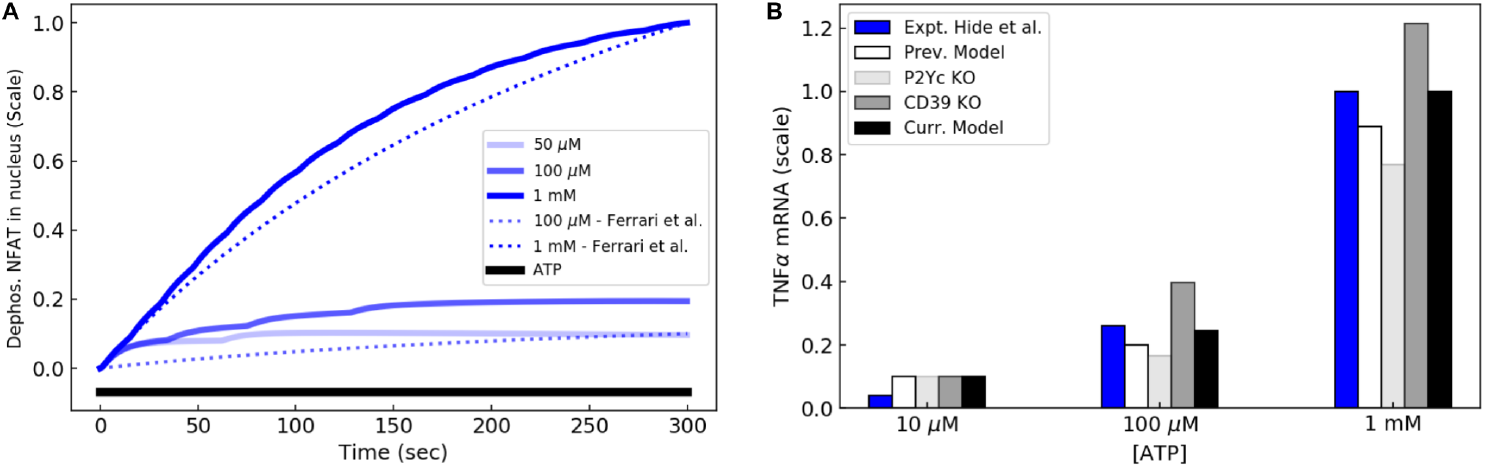
Predictions of dephosphorylated NFAT in nucleus over time (A) and TNF*α* mRNA with various computation configurations and comparison to the previously developed model(9) with respect to amplitude of stimulation (5). All simulations in B) were performed for 5 minutes. The plot is in the unit of scale, whose basis is the maximum increment measured by the current model.

**Figure S8:**
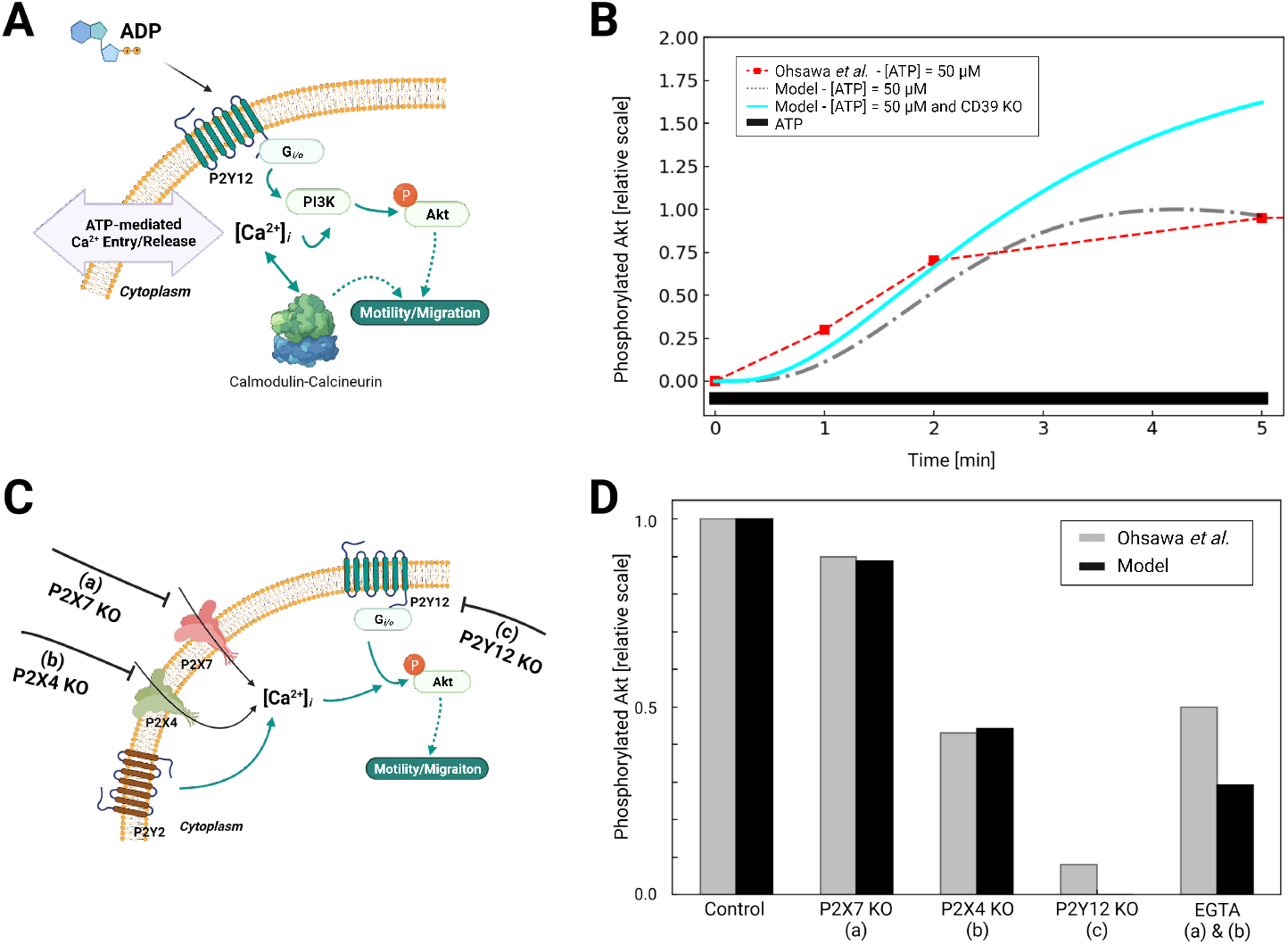
A) Schematic of Akt phosphorylation via P2Y12- and P2Y-mediated Ca^2+^ signaling pathways in response to ATP. B) Predicted *p*Akt expression as a function of time in response to 50 uM ATP applied for 5 minutes with and without CD39 expression. Experimental data for 50 uM ATP from Ohsawa *et al* are presented in red. C) Schematic of P2Y12- and Ca-mediated migration in response to ATP, assuming control, P2X7 knockout (KO) (a), P2X4 KO (b) and P2Y12 KO (c) conditions. D) Predicted *p*Akt expression (black) versus experimental measurements by Ohsawa *et al* under control and a-c conditions. .

**Figure S9:**
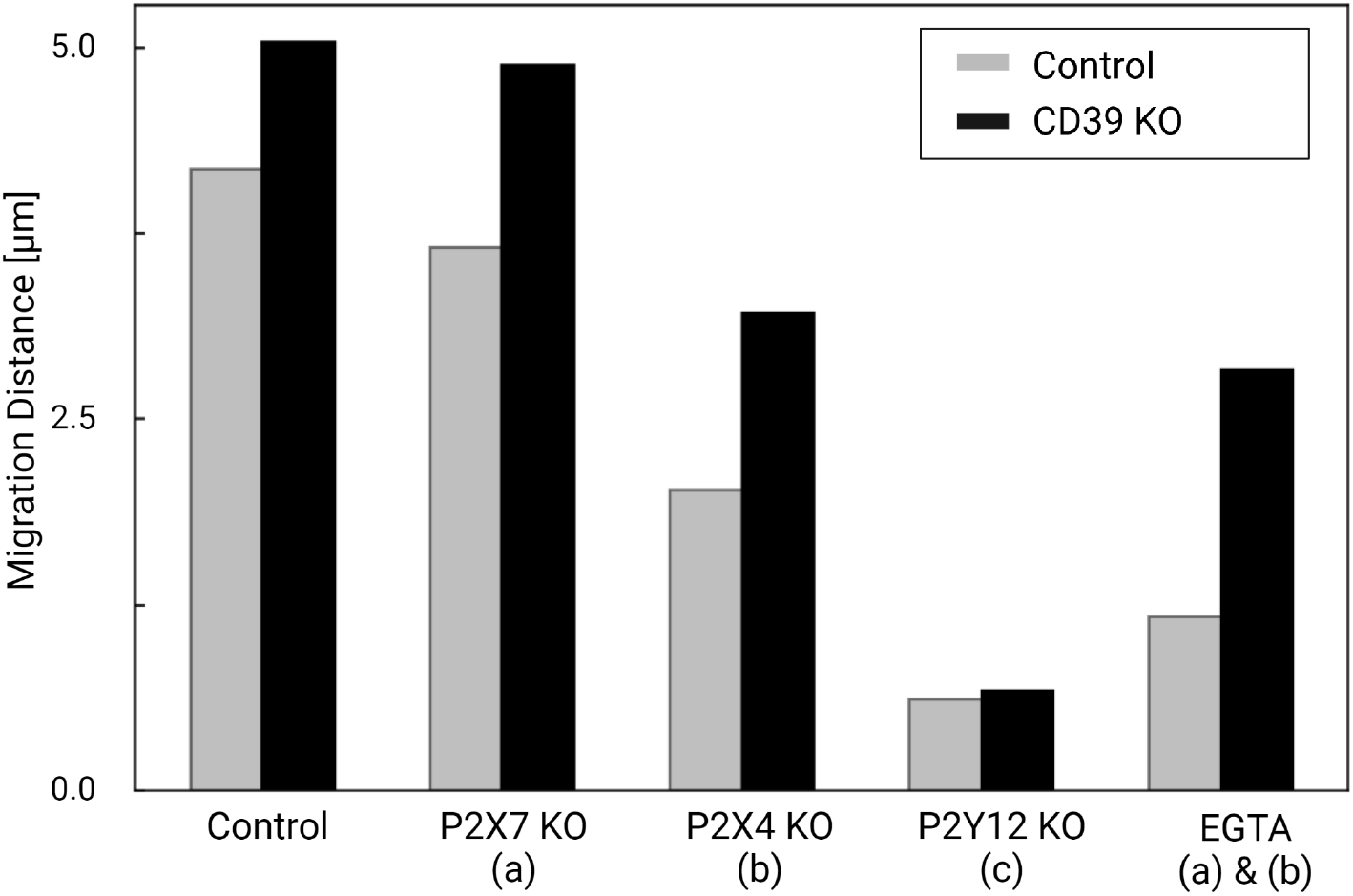
Migration comparison between WT and CD39 KO models.

**Figure S10:**
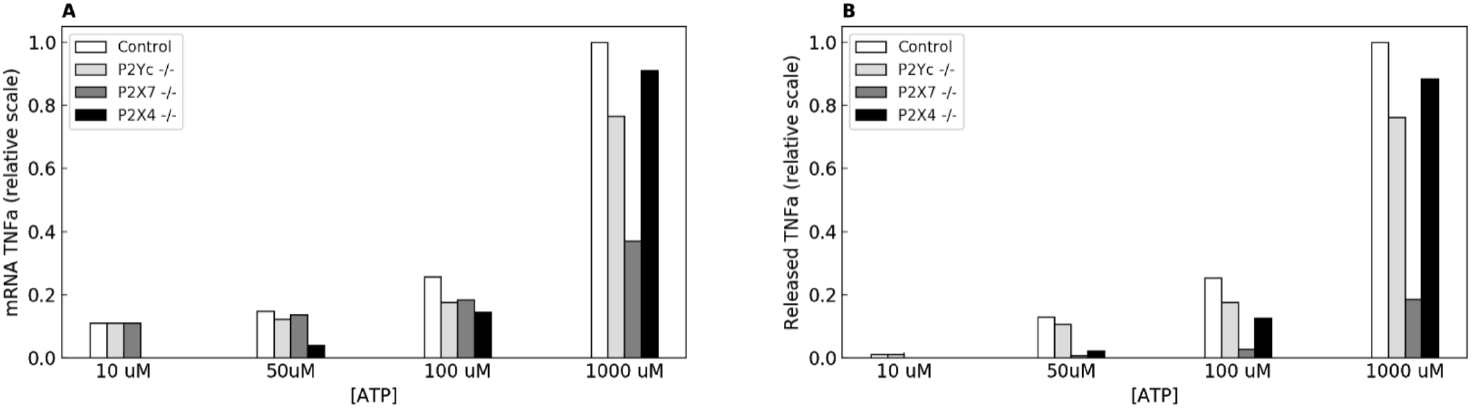
Prediction of TNF*α* mRNA expression(A), and TNF*α* release(B) with conditions of *P*2*Y* −/−, *P*2*X*7 −/−, and *P*2*X*4 −/−. .

**Figure S11:**
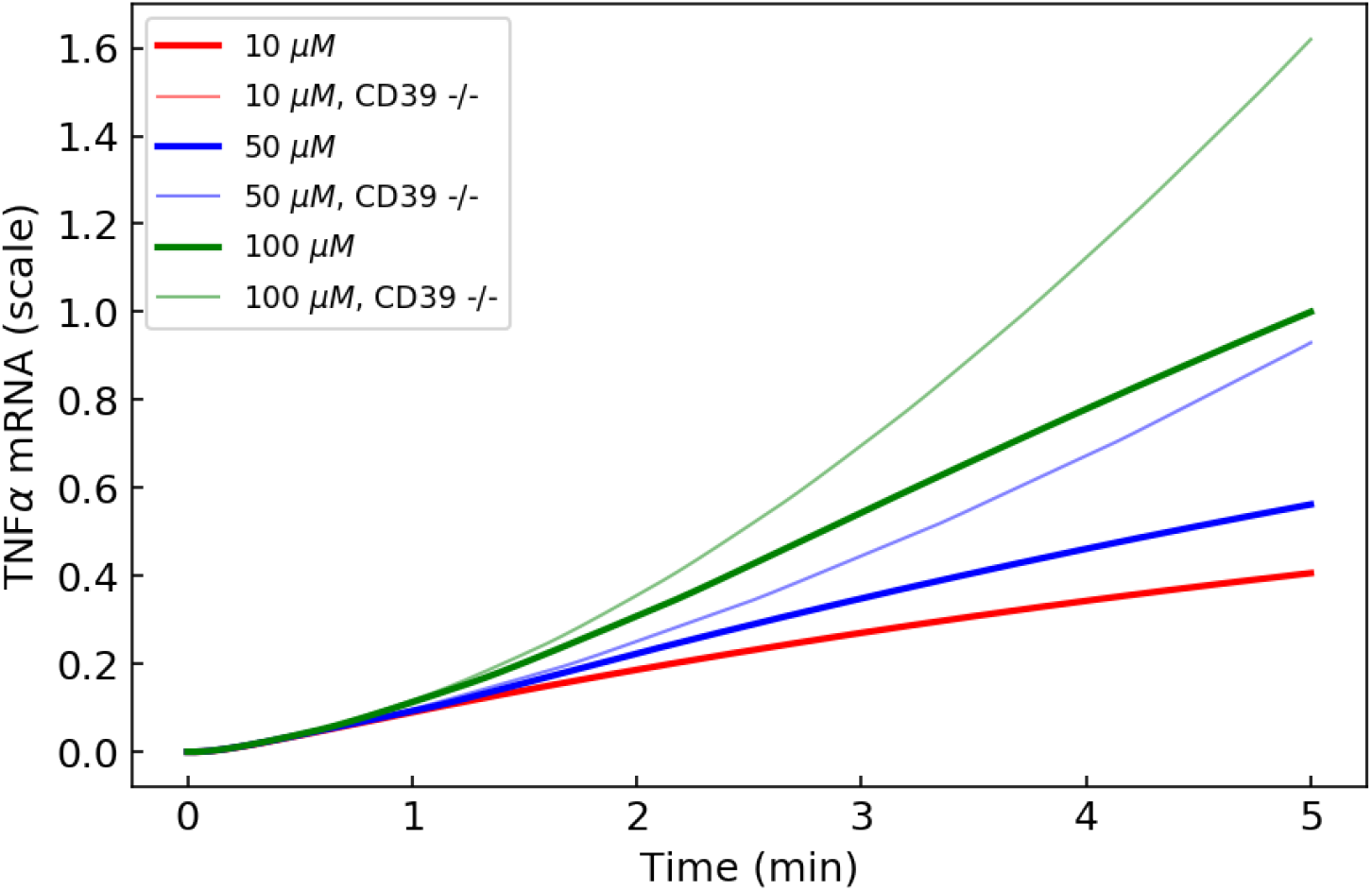
TNF*α* prediction with low ATP and with/without CD39.

**Figure S12:**
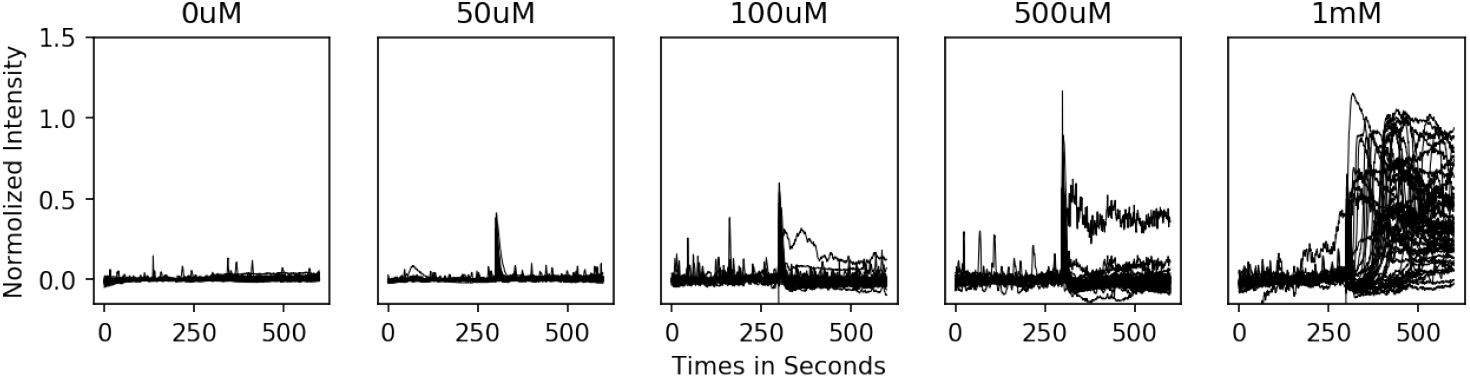
Traces of the fluorescence to measure the ATP-mediated Ca^2+^ transients in BV2 cells.

**Figure S13:**
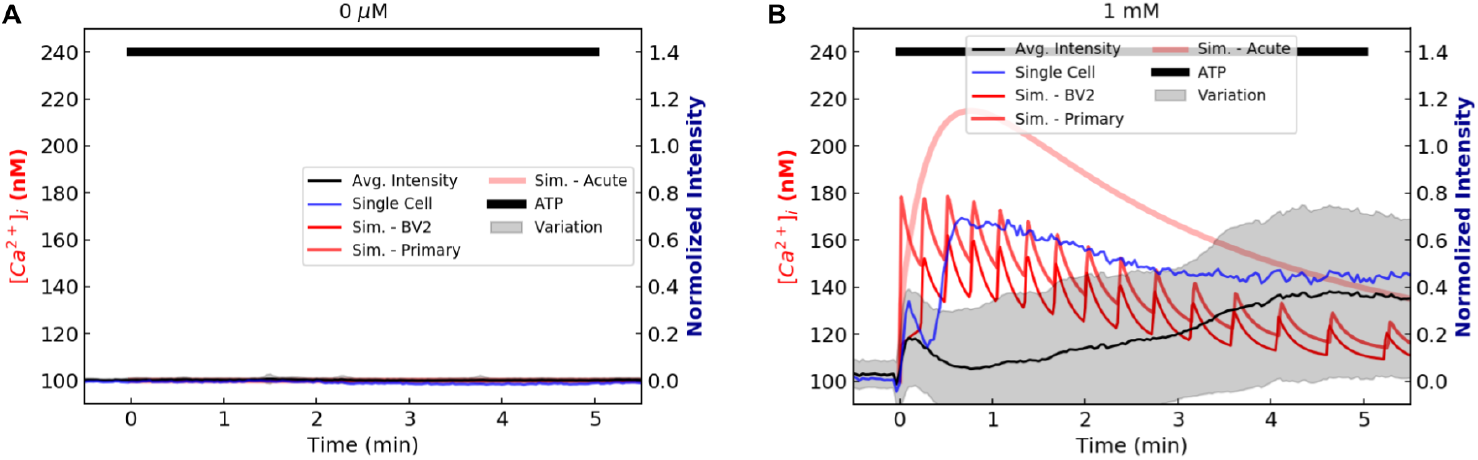
Predictions and experimental data of Ca^2+^ transients in BV2 with 0 μM and 1 mM ATP. .

### S.5 Results

#### S.5.1 E-NTPDase (CD39): ATP decomposition to AMP

Ectonucleoside triphosphate diphosphohydrolase-1 (E-NTPDase) also known as CD39 is reported to be present in the surface of microglial plasma-membrane and play an important role in microglial migration by balancing ATP and adenosine molecules (25). Using the generic ATP decomposition into ADP and AMP by CD39 introduced in the (47), we have added the ATP/ADP/AMP correlation to the previous model to properly differentiate the ATP-mediated and ADP-mediated activation of purinergic receptors. Therefore, the stimulant for P2X and P2Y-class that mediates Ca^2+^ fluctuations is ATP whereas P2Y12 is specifically stimulated by ADP.

#### S.5.2 Migration

In order to verify that migration occurs within the time intervals of which ATP-induced Ca^2+^ transients were measured, we report changes in above-threshold cell areas before and after ATP treatment.

### S.6 TNF*α* production mediated by purinergic receptors

#### S.6.1 Validation of TNF*α* mRNA data

A primary consequence of ATP activation in microglia is the production of TNFa (33). Our previous model of ATP-triggered TNF*α* production in microglia we validated in (9) was limited to *P*2*X*4 and *P*2*X*7. We therefore confirm that the introduction of *P*2*Y* and ectonucleotidase activity contributions maintains TNFa mRNA production rates consistent with experiment. We first evaluate the level of activated (dephosphorylated) NFAT, which is a prominent transcription factor driving TNFa mRNA synthesis In Fig. S7A we demonstrate an ATP dose-dependent increase in activated NFAT that remains in close agreement with experimental data from *et al*. In Fig. S7B, we illustrate the corresponding predicted TNFa levels. The data used for this validation was based on TNF*α* release measured by Iketa *et al* which was reported at under 1 mM ATP treatment. For validation, we assumed that mRNA production was proportional to TNF release, and thus we rescaled the TNFa release data at 1 hour/1mM ATP to 5 minutes for 10 μM-1 mM ATP. The predicted TNF*α* mRNA levels for control remain in close agreement with comparable conditions imposed in Hide *et al* (33). The comparable results between the previous model (white) and current model (black) confirm that the P2Y/ENT implementations remain compatible with our previous validation data. This also suggests that the contributions of *P*2*Y* do not significantly contribute to TNFa mRNA responses, as *P*2*Y* knockdown in our model generally reduces the predicted mRNA levels by less than 10%. We also present data for which ectonucleotidase activity is neglected, which demonstrate amplified TNFa mRNA production. The data indicate that P2Y receptors do not significantly impact TNF*α* production, owing to CD39 activity that attenuates their responses.

### S.7 Relative contribution of P 2X vs. P 2Y to cytokine production and migration

The preceding section walked through the procedure and validity of modeling microglial responses to ATP stimulation. We note that a subtype of P2 receptors may mediate one or more pathways toward the production of pro-inflammatory cytokines, such as TNF*α*, instead of the initiation of chemotaxis or vice versa. For instance, the *P*2*X*-mediated Ca^2+^-entry effectively promotes intracellular Ca^2+^-dependent signaling that leads to TNF*α* secretion. On the other hand, the activation of metabotropic receptors trigger various pathways whose outcomes include Ca^2+^ fluctuations and structural responses such as migration. To quantitatively analyze the functionality of specific P2 receptors in response to ATP, we carried out a series of simulations that selectively inhibit individual receptors. We additionally performed simulations with varying ATP concentrations to maximize the differential activation of P2 receptors.

Fig. 5 showcases the ATP-mediated mRNA transcription and secretion of TNF*α* and corresponding metrics of chemotaxis for each calculation configuration. We observed the expected absence of increase in basal Ca^2+^ levels in the *P*2*X*7 KO simulation (Fig. 4). This resulted in a noticeable reduction in the TNF*α* mRNA transcription and release of matured cytokines compared to controls. On the contrary, in the lower range of ATP (up to 100 μM), the activation of *P*2*X*4 receptors promote Ca^2+^ fluctuations sufficient for mRNA transcription. It is apparent that the differential degrees of ionotropic receptor activation throughout the tested ATP spectrum directly influence the Ca^2+^-entry, and it is well reflected on the obtained mRNA data. However, the inhibition of metabotropic receptors shows a relatively trivial change in the outcomes. The difference between inhibition of *P*2*Y* 12 and *P*2*Y*-class receptors is, though, that *P*2*Y*-class activation triggers ER Ca^2+^ release, whereas *P*2*Y* 12-mediated secondary messenger elicits neither Ca^2+^ fluctuation nor TNF*α*-associated pathways.

The predicted secretion of TNF*α* is consistent with the preceding mRNA data except for the data of *P*2*X*7 KO setup. Suzuki *et al* (75) claimed that TNF*α* release is correlated with the activation of *P*2*X*7, the inhibition of which by brilliant blue G (BBG) resulted in up to 50% reduction in released TNF*α* from microglia after 15-minute prolonged stimulation with 1 mM ATP. The simulated TNF*α* release at the same ATP condition slightly overshoots the degree of reduction (nearly 60%) in the production/release of TNF*α*.

Simulations of chemotaxis display the relatively extensive dependency on *P*2*Y* 12 receptor activation (Fig. 5D). Silencing *P*2*Y* 12 receptors results in a substantial truncation of migration throughout the simulated range of ATP (more than 90% of reduction up to 1 μM, and 50% with 1 μMATP). Like with TNF*α* synthesis, the magnitude of reduction in migration induced by *P*2*X* receptors varies as ATP concentration varies due to their varying sensitivities to ATP. Additionally, their deletion does not cause as severe a reduction in migration as knocking out *P*2*Y* 12 receptors. Overall, our prediction indicates the substantial involvement of *P*2*X*7 receptors in the cytokine production; the *P*2*Y* 12 activation is crucial for invoking pathways associated with microglial chemotaxis.

